# Cellular code for mnemonic pattern separation in the human hippocampus is revealed by false memories

**DOI:** 10.64898/2026.09.18.752786

**Authors:** Natalia Kurilenko, Clayton Mosher, Sophia Cheng, Yousef Salimpour, Jonathan Daume, Chrystal M. Reed, William S. Anderson, Taufik A. Valiante, Adam N. Mamelak, Ueli Rutishauser

## Abstract

Theoretical models propose that pattern separation, a computation thought to be implemented by the hippocampus, allows us to differentiate between familiar and novel items. However, whether pattern separation operates in the human hippocampus remains contested, with no established single-cell correlate. We recorded the single neuron activity of 3506 neurons in the human brain while 97 patients performed a recognition memory task in which novel images similar to previously seen images led to false memories and associated behavioral errors. We identified two kinds of memory selective neurons distributed across the brain: those responding differently to falsely recognized novel and correctly recognized familiar images in a manner compatible with pattern separation, and the other signaling the subject’s choice. At the population level, these cells predicted mnemonic ground truth in the hippocampus and the decision in the pre-supplementary motor area, illustrating the progression from mnemonic signals to decisions. Removing pattern separation-signaling cells abolished the continuous memory strength gradient present in the hippocampus, suggesting a role of these cells in separating memories of different strength. These results establish a single cell correlate for mnemonic pattern separation in the human hippocampus and show its behavioral relevance in episodic memory.

## Introduction

Humans can form large numbers of detailed memories of individual experiences^1–4^, an ability that allows us to plan, learn, and make decisions. Despite many of our experiences taking place in highly similar environments (such as our home or office), many are encoded as distinct memories. A major open question is how the brain achieves this feat. Solving this problem requires resolving a trade-off between two seemingly incompatible aims^5,6^. On the one hand, a familiar item should be recognized in many different contexts, requiring generalization and invariance. On the other hand, novel items should be distinguished from stored memories even if very similar, requiring fine mnemonic discrimination. Critically, this discrimination must often occur after a single exposure (such as in episodic memory). These complementary processes are often referred to as pattern completion and separation, respectively^6^. Accordingly, a major focus of the field has been on identifying signatures of these two processes in the hippocampus (HPC), a region essential for memory formation. However, despite an extensive literature^6–10^, it remains unknown whether pattern separation plays a role at all in human episodic memory^11–15^. A key reason for this large gap in knowledge is that no single neuron correlate of pattern separation has been identified in humans. Here, we identify such a correlate for memory strength signals.

Neural evidence for a role of pattern separation in human memory comes principally from non-invasive neuroimaging using fMRI. At the BOLD-fMRI level^9,10^, activity in the HPC exhibits an absence of repetition suppression to “lure” stimuli, which are novel stimuli that are similar to items previously seen. This pattern has widely been taken as evidence for successful discrimination between very similar inputs and thus reflective of the process of pattern separation^16–18^. However, several distinct theorized mechanisms — including conjunctive coding and match-mismatch or novelty signaling — could give rise to this very pattern of BOLD-fMRI signals^11,19^, making this approach unsuitable to mediate between the different theories. In addition, in these studies, it has not been shown that this signature of pattern separation is related to mnemonic processes because participants’ memory for the presented stimuli was not tested^16–18^.

At the level of single neurons, in contrast, most work to date suggests that pattern separation is not present in the human brain. For example, recordings from cells in the human HPC show that concept neurons encode stimulus identity in an abstract manner, with no sensitivity to even relatively large differences as long as the stimulus is from the same concept or visual category^20–23^. This pattern of response is the opposite of what would be expected if the activity of these cells is the result of pattern separation. This is because the highly invariant firing leads to overlapping representations for similar yet distinct inputs, making them indistinguishable. These findings support the conclusion that pattern separation plays no role in human episodic memory^11^. Instead, it has been proposed that episodic memories are encoded by partially overlapping assemblies of concept cells^11,24^, a framework in which pattern separation plays no role. This proposal sparked extensive debate^11–15^ that remains unresolved. Here, by identifying a single-neuron correlate, we resolve this debate by showing evidence for pattern separation in the human hippocampus.

In rodents, on the other hand, evidence for pattern separation is strong in the domain of spatial coding^25–31^. The major finding is that hippocampal place cells change their activity (‘remap’) in response to even subtle changes in the environment, which results in a decorrelated code for the two similar but distinct enviornments as one would expect from pattern separation^26,27,31^. Theoretically, an explanation for this finding has been that it is the result of conjunctive coding^6,7^, a kind of coding that often results from a high fan-out ratio. For example, in the hippocampus, the sparse projections from the entorhinal cortex to the hippocampal dentate gyrus (DG) result in individual DG neurons only being activated by unique combinations (the conjunction) of input features. In this coding scheme, different populations of DG neurons are recruited for two similar stimuli (here, two different locations), since even small changes in the input will alter which combination of features is present.

The original proposers of pattern separation suggested that its principal role is to enable an organism to distinguish between familiar and similar to them novel stimuli^5–7,9^. Behaviorally, this ability is impaired by lesions in parts of the medial temporal lobe that are, based on rodent studies, thought to be important for neuronal pattern separation. In particular, patients with damage to the DG show impaired discrimination between previously seen and highly similar images—thus lacking the behavioral hallmark of pattern separation^32,33^. However, this aspect of pattern separation was not examined in either the rodent studies nor the concept cell studies in humans, which instead both tested pattern separation between representations of the content of an experience (spatial and concepts present, respectively). As a result, these studies did not address the aspect of pattern separation most relevant for memory: the distinction between novel and familiar stimuli. In contrast, this is precisely the comparison tested in the relevant fMRI studies^16,17^, which do report evidence consistent with pattern separation.

To fill this gap, we examined whether memory strength signals in the human temporal lobe^23^ exhibit signatures of pattern separation. We did so by recording single-neuron activity in epilepsy patients while they performed a recognition memory task images with varying similarity to each other.

## Results

### Task and behavior

97 patients (133 sessions) performed a recognition memory task with pictures as stimuli. First, during a learning phase, patients viewed a series of novel images. Then, during a recognition phase, the same images were presented again, intermixed with novel images (Fig. 1a). For each image patients indicated whether they had seen it before (“old”) or not (“new”), and reported ther confidence in that judgment (Fig. 1a). The task has 3 variants that are identical in design but employed different stimuli.

**Figure 1.**
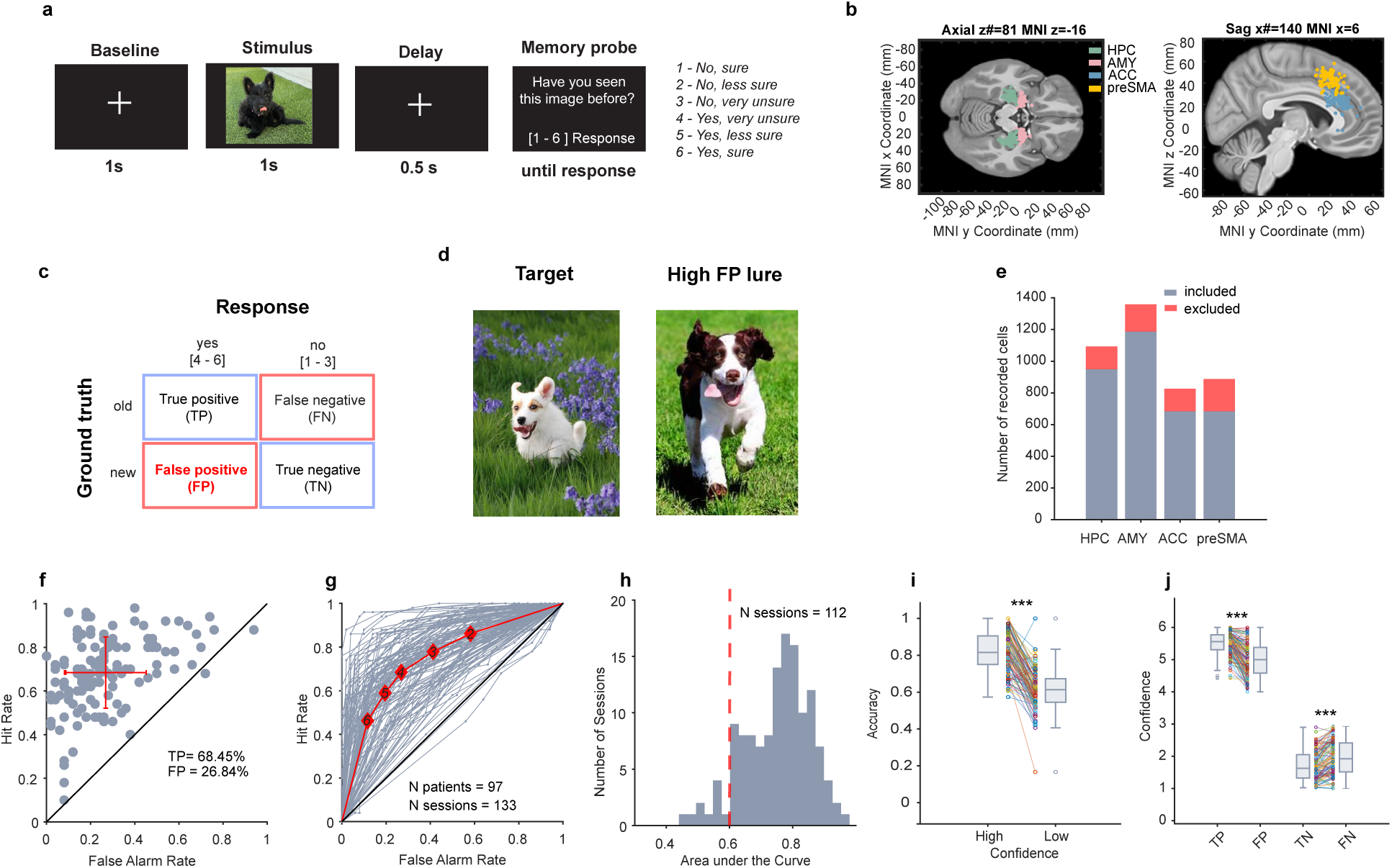
Task, behavior, and summary of recorded cells. **a**, Example recognition trial. **b**, Electrode locations displayed on an Atlas brain. Each dot represents a microwire bundle, colored by recording area. White matter or off-target coordinates are due to the merge onto the template brain. **c**, Possible trial types: False positives (FP highlighted with red) result when a subject categorizes a ‘new’ image as ‘old’ (Yes response). **d**, Left: Example target image (old) that was studied. Right: visually similar lure image that was not studied (new), but which was classified as old (an FP) with high probability (FP frequency = 54%). **e**, Number of isolated single units (grey, n total across all brain areas = 3506) and number of units excluded based on behavior (red, n=660). **(f-j).** Behavior. **f**, Recognition memory accuracy. Each point is one session (n = 133 across 97 patients), red is the mean ± s.d. (**g-f)**, Accuracy as a function of confidence for individual sessions (gray) and average (red). Each data point is a different confidence level, with 6 being the highest confidence for the old trials. **h**, AUC under the ROC curve shown in (**g**). The dashed red line is one of the inclusion criteria (AUC>=0.6, included n = 112, 84 patients). **i**, Accuracy was significantly higher in high vs. low-confidence trials (p=1.24×10-31, t(110)=16.55, paired t test). Each line is a different session. **j**, Incorrect responses (‘yes’ or ‘no’) had significantly lower confidence ratings compared to correct responses (TP vs FP: t(110)=13.41, p= 7.05×10-25, paired t test; TN vs FN: t(111)=-8.72, p= 3.11×10-14, paired t test. Note that negative t statistics are due to higher confidence rating corresponding to lower confidence for response ‘no’, see panel a). **\*\*\*** p<0.001

Patients correctly identified 68.5±16.3% (Mean±s.d) of the old images (true positive, TP or hit rate; Fig. 1f, y axis). At the same time, they incorrectly reported as “old” 26.8±18.3% (Mean±s.d) of the novel images, trials which we refer to as false positives (FP, Fig. 1f, x axis). Recognition accuracy increased as a function of confidence (Fig. 1g, i-j)^23^. To quantify the relationship between recognition accuracy and confidence, we computed a receiver operating characteristic (ROC) curve for each session (Fig. 1g). Trials with different confidence ratings fell at distinct points along the ROC curve, reflecting systematic differences in performance across confidence levels (Fig. 1g). The average area under the ROC curve (AUC) was 0.75±0.1 (Fig. 1h). Pairwise comparisons across sessions showed that trials reported with high confidence (1 for ‘new’, 6 for ‘old’) had significantly higher accuracy than those reported with low confidence (2–3 for ‘new’, 4–5 for ‘old’; 0.82±0.11 vs. 0.61±0.11 (Mean±s.d), p=1.24×10^−31^, paired t-test, Fig. 1i). Incorrect trials had significantly lower confidence ratings than their same-response correct counterparts (TP vs FP: 5.5±0.35 vs. 5±0.52, p= 7.05×10^−25^; true negative (TN) vs false negative (FN): 1.69±0.45 vs. 1.95±0.53, p= 3.11×10^−14^, paired t-test, all Mean±s.d; Fig. 1j). These results support the validity of the subjective confidence ratings in both correct and error trials.

### Perceptual similarity drives false alarms in recognition memory

Patients made two types of mistakes (Fig. 1c, red boxes): they either forget an image they have seen before (FN) or they identified a new image as old (FP). Here, our focus is on FP errors because it is during that type of trial where pattern separation putatively failed^34^. We hypothesized that FPs are the result of new images being similar to previously studied items (Fig. 1d shows an example), leading them to be rated as old for this reason. If so, FPs are a result of the failure of separating two patterns – that stored in memory and the image shown. We tested this hypothesis by defining and validating similarity metrics for pairs of the visual stimuli used, followed by testing whether the similarity metrics predicted the likelihood of making a FP error for a given image (Fig. S2a-h).

We first tested whether pre-trained deep networks can predict how similar a pair of images is perceived by human subjects. We used (i) AlexNet and ResNet50, two convolutional neural networks (CNNs) trained for object categorization, (ii) DINOv2, a self-supervised vision transformer (ViT) that extracts general-purpose visual features, and (iii) CLIP, a ViT-based model trained with natural language supervision to align text and images. As a control, we included models that only use (i) low-level visual features, and (ii) a semantic ‘categories’ model, which was built by assigning a dissimilarity of 0 to image pairs within the same semantic category and 1 to pairs from different categories. Together, these models span different architectures, training strategies, and feature domains.

To assess how well these models capture human judgments of similarity for the specific stimuli used in our task, we also asked human subjects to rate the similarity of a subset of images (Fig. S2i; 25 images with five images in each of five categories; see Methods). We then constructed representational dissimilarity matrices (RDMs) for all models (Fig. S2j) and compared the resulting RDMs with that of the human subjects (see Methods^35^). All models, except for the pixel-wise model, performed significantly better than chance (one-sided t-test, Bonferroni-corrected α = 0.002 for 6 models), with the highest performing models being CLIP and DINOv2 (Fig. S2k). These two models outperformed the category model (p=0.0001 and p=7.7×10^−12^ respectively; FDR-corrected pairwise t-test for 15 model-pairs comparisons), suggesting that human similarity judgments are better explained by models that capture rich visual and semantic features, rather than by categorical labels alone.

Finally, we tested whether the derived similarity metrics predicted FP rates (Fig. S2d,h). To do so, for each new image *i* shown during retrieval, we computed the probability of incorrectly recognizing this image as “old” (the false alarm rate) across all patients who have seen this image (n=41 for task variant 1, n=16 for variant 2, and n=53 for variant 3). Deep neural networks (DNN)-derived similarity metrics between a given new image and all previously shown images significantly predicted false alarm rates (Fig. 2a shows the ‘CLIP’ model; Spearman r=0.4, p=0.009; r=0.7, p=4.9×10^−10^; r=0.5, p=0.0006 for variants 1, 2, and 3, respectively). This was also true for human-derived similarity judgments, which predicted FPs with similar (if not lower) accuracy (Fig. 2b; Spearman r=0.3, p=0.02). This data shows that DNN-derived pairwise similarities are good predictors of FP rates.

**Figure 2.**
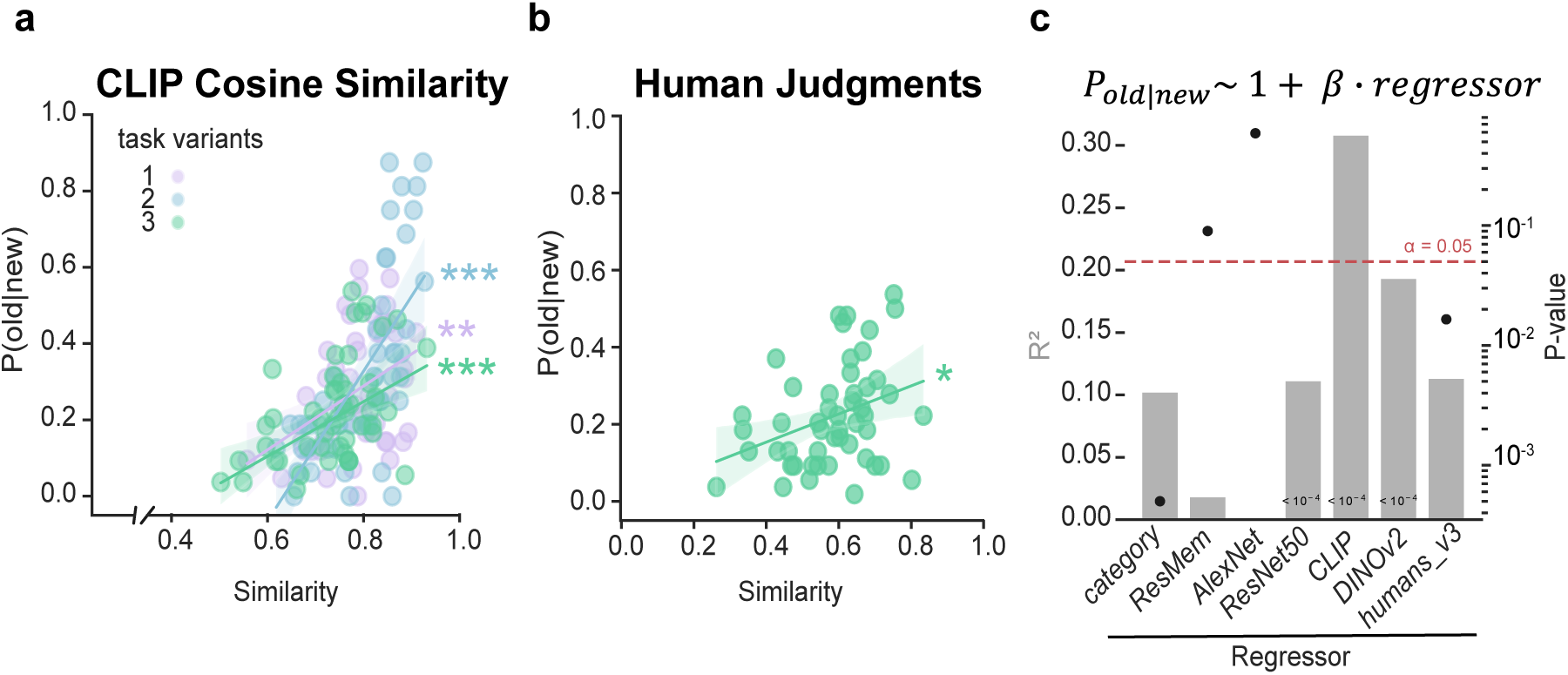
False alarm probability is predicted by perceptual similarity between image pairs. **a,** Correlation between DNN ‘CLIP’ based similarity (maximum similarity between each ‘new’ image presented during the recognition part of the experiment (Fig.1a) and all learning images) and false positive (FP) rate calculated across all patients, color-coded by task variants. For all 3 task variants, image similarity was significantly correlated with FP rate (r=0.4, p=0.009; r=0.7, p=4.9×10–10; r=0.5, p=0.0006 for variants 1, 2, and 3, respectively, Spearman correlation). **b,** Correlation between human perceptual similarity (maximum similarity between each ‘new’ image and all learning images from the same semantic category) and false positive rate (r=0.3, p=0.02, Spearman correlation); judgments were collected only for task variant 3. Each dot in **a** and **b** is a ‘new’ image. **c,** Statistical comparison of repeating analysis shown in panels (a,b) for different models. Visual similarity computed with different features (ResNet50, CLIP, DINOv2, human judgments) explained a significant amount of variance in false positive rates, while image memorability (ResMem) did not. The left y-axis corresponds to R-squared (grey bars), and the right y-axis corresponds to regression p-value, with individual values depicted as black dots on the plot. The red dashed line marks the alpha level of 0.05 used to denote statistical significance. * p<0.05, ** p<0.01, *** p<0.001.

False alarms can also arise due to factors other than visual similarity^36^, including an image’s intrinsic sense of familiarity^37^ and category membership^4^. As a control for these two factors, we also included in our analysis semantic category and memorability scores quantified with ResMem, a neural network trained to predict image memorability^38^. Comparing the amount of explained variance between the CLIP and DINOv2 derived metrics with these two control models shows that neither explained nearly as much variance. Indeed, image memorability (ResMem) did not account for a significant proportion of the variance in FP probabilities (Fig. 2c, r2=0.02, p=0.1). Together, these findings support our hypothesis that false alarms in our patients during our task arise, at least in part, from new images being similar to previously studied ones. This finding suggests that in our task, false alarms are due to failures in pattern separation.

## Recordings

While patients performed the task, we recorded single-neuron activity bilaterally from HPC (n=1093), amygdala (AMY, n=1359), dorsal anterior cingulate cortex (ACC, n=826), and pre-supplementary motor area (preSMA, n=888; Fig. 1b,e). Out of all sessions recorded, we included only the sessions with sufficient behavioral accuracy (patient-level AUC > 0.6, red dashed line in Fig. 1h) and with responses spanning both high- and low-confidence ratings (n=112 sessions satisfied these criteria, see Methods). Across all included sessions, we recorded 950, 1187, 684, and 685 cells in HPC, AMY, ACC, preSMA, respectively (3506 total, Fig. 1e).

### Memory-selective cells carry a ground-truth memory signal even when behavior is incorrect

We next examine the characteristics of neuronal responses associated with FPs. We did so by examining memory selective (MS) cells, which respond differently to novel and familiar stimuli in a memory strength-dependent manner^23^.

We identified MS cells by contrasting firing rates during stimulus presentation between correct new and old trials (Fig. 3a,b shows examples; see Methods; correct here refers to the accuracy of the patients response in that trial). To avoid confounds related to stimulus category, we also selected cells that differentiated between categories using a one-way ANOVA (1 × 5) on firing rates across trials containing stimuli from each category. A significant proportion of cells qualified as MS cells in all four recording sites (Fig. 3c). Out of all cells, 10% (94/950) in the HPC, 9% (102/1187) in the AMY, 9% in the ACC (60/684), and 16% (113/685) in the preSMA (chance level is 5%; see Fig. S3a-e for absolute numbers) qualified as MS cells. On the other hand, MS cells that were also selective for visual categories (visually-selective, VS cells) were rare (<2% across all the brain areas, Fig.3c). The responses of MS cells were modulated by memory strength in all brain areas except in the ACC: their signal was stronger in high confidence compared to low confidence trials (Fig. 3d, HPC AUC 0.65±0.01 vs. 0.6±0.02, p=0.006, AMY 0.66±0.01 vs. 0.61±0.01, p=0.0009, ACC 0.66±0.01 vs. 0.62±0.03, p=0.1, preSMA 0.68±0.01 vs. 0.64±0.01, p=0.03, paired t-test; Fig. 3e for AUC comparison separately for novelty and familiarity preferring cells across all areas).

**Figure 3.**
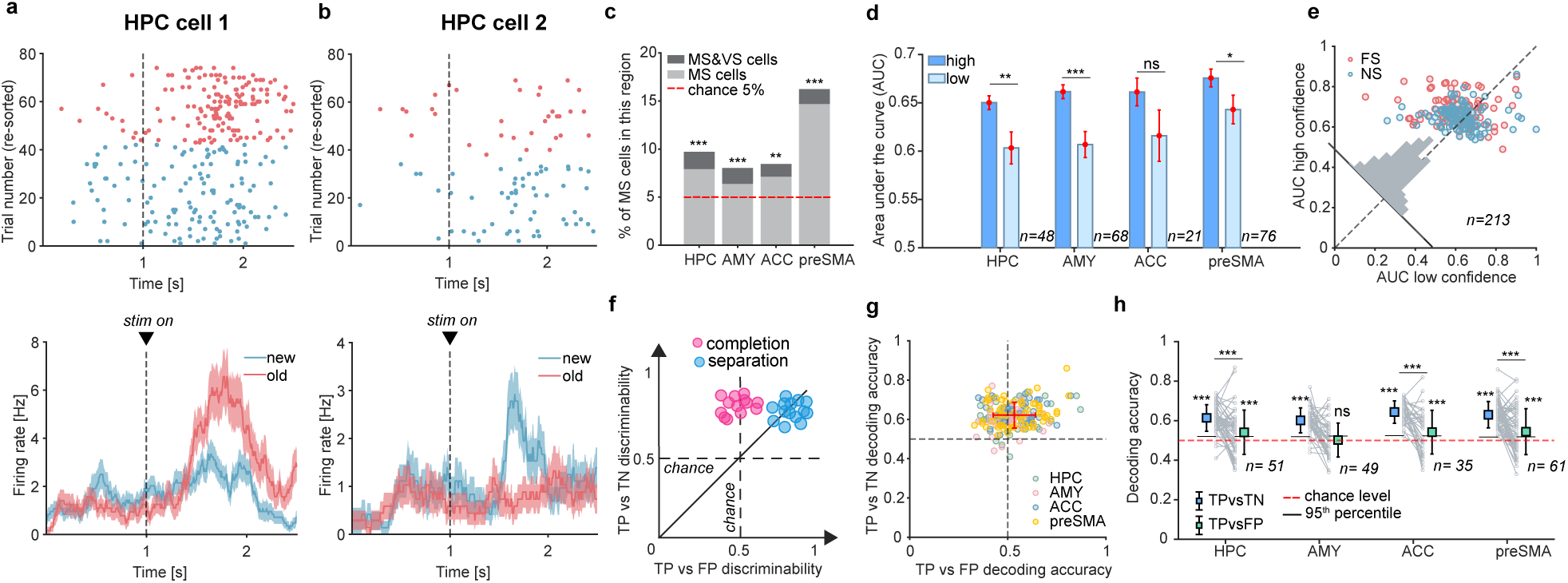
Single-neuron signatures of recognition memory: Memory selective cells. **(a-b)** Example memory selective cells. Stimulus onset is at t=1. **a,** Example hippocampal neuron that increased its firing rate most for correctly recognized old vs. new stimuli. **b,** Different example cell, which increases its firing rate for correctly recognized new vs. old stimuli. **c,** Proportion of all recorded cells that qualified as memory selective (MS) cells. Dark grey color indicates MS cells that in addition also qualified as VS cells. The proportion of MS cells is larger than expected by chance (red line) in all areas (HPC, p=0.001, AMY, p=0.001, ACC, p=0.002, preSMA, p=0.001) **d, e** MS cell signal is stronger in high compared to low confidence trials. d, Pairwise comparison of MS cells AUC values for high and low confidence by brain areas (p-values are paired t-test). MS cells in all brain areas except for ACC (p=0.1) had significantly higher AUC for high confidence trials than low confidence (HPC p=0.006, AMY p=0.0009, preSMA p=0.03). Only behavioral sessions where patients could differentiate high and low confidence were included in the d and e analysis (69 sessions across 53 patients). **e,** Same as **d**, but plotted as a scatter plot separately for novelty- (blue dots) and familiarity- (red dots) preferring cells across all 4 brain areas. Note that for both types of MS cell AUC for high confidence trials is higher than for low confidence. **f,** Illustration of hypothesis: the activity of neurons that employ pattern separation (blue dots) reliably distinguishes between true old (TP) and false old (FP) stimuli, despite the response in either case being ‘old’. Conversely, neurons exhibiting pattern completion–like activity (magenta dots) would respond to FP trials in the same or similar manner as they do to TP trials. Dots show hypothetical simulated data. **(g-h)** Data to test the hypothesis shown in (f). Across all recorded MS cells, activity differed significantly between TP and FP trials, indicating the presence of a memory rather than a decision signal. **g,** Cross-validated single-cell decoding performance for neurons that were memory-selective in at least 75% of randomly selected trials, color-coded by brain area. Red is the mean ± s.d. For many MS cells, decoding performance of TP vs FP exceeded the chance level. **h,** Pairwise comparison of TP vs TN and TP vs FP decoding performances for cells from g. In HPC, ACC, and preSMA, both TP vs TN and TP vs FP were decodable significantly above chance (p=0.001), with TP vs TN decoding being significantly higher than TP vs FP (HPC p=7.5×10-5, ACC p=7.6×10-5, preSMA p=7.4×10-6, signrank test). In AMY, only TP vs TN was significantly decodable (p=0.001, TP vs FP p=0.4). * p<0.05, ** p<0.01, *** p<0.001.

How did MS cells respond during FP trials? We hypothesized that if the response of MS cells is reflective of pattern separation, their responses should distinguish between true old (TP) and FP trials, despite the behavioral response being the same (‘old’) in both cases (Fig. 3f, blue). Given the prominence pattern separation theory provides to the hippocampus^5–7^, we further predicted that this difference is particularly strong in the hippocampus. In contrast, if the response of MS cells during FP trials is the same as that during TP trials (Fig. 3f, magenta), the cells would signal a decision about the memory instead, and would thus be reflective of pattern completion. To avoid confounds due to simultaneous visual category tuning, we excluded the small number of MS cells that were also visually tuned for all subsequent analysis (see Methods; doing this did not change the results).

To assess the response of MS cells during FP trials, we trained a binary single-cell decoder to classify stimuli presented during correct trials as new or old. If, for left out trials in the training set, new vs. old decoding accuracy was above chance, a cell was classified as an MS cell. We then assessed decoding performance on left-out trials—unseen during MS cell selection—for either TP vs. TN (correct old vs. correct new) or TP vs. FP (correct old vs. incorrect new) discriminations. This approach allowed us to compare decoding performance for both correct and false alarm trials across brain areas (we only used cells that were classified as MS cells in at least 75% of the train/test splits). The single cell decoders reliably decoded memory from held out correct trials across brain areas (Fig. 3g-h, y axis; HPC decoding performance 0.61±0.07, AMY 0.60 ±0.06, ACC 0.64±0.06, preSMA 0.63±0.07, p=0.001; Chance is 0.5). Turning to the error trials, decoding of TP vs. FP trials was, on average across all MS cells, above chance in the HPC (0.54±0.11, p=0.001) as well as ACC and preSMA (0.54±0.11 and 0.55±0.12, respectively; p=0.001 for both). In all three sites where ground truth of FP trials was decodable, TP vs. TN decoding was significantly more accurate compared to TP vs. FP (Fig. 3h; HPC p=7.5×10^−5^, ACC p=7.6×10^−5^, preSMA p=7.4×10^−6^, signrank test), underscoring behavioral relevance. This finding is compatible with the presence of a pattern-separation related signal in MS cells in the HPC, ACC, and preSMA as a group.

### Single-neuron signature of pattern separation: two types of MS cells across MTL and MFC

Were there different types of MS cells, with some reflecting pattern completion and others pattern separation as hypothesized (Fig. 3f)? To answer this question, we tested the ability of individual MS cells to discriminate between trials with different ground truth but the same behavioral choice (TP vs. FP), or trials with the same ground truth but different choices (TN vs. FP).

Fig. 2 shows that FPs are predicted by the similarity of stimuli with those previously studied (TP trials). If the response of a cell discriminates between these two types of trials (TP and FP) despite the same behavioral outcome (“old”), but not between TN vs. FP trials, we call this cell a ‘pattern separation’ cell (Fig. 4a, blue dots). In contrast, if an MS cell signals the choice, its response should discriminate between trials with different choices despite the same ground-truth (TN vs. FP), but not trials with the same choices and different ground truth (TP vs. FP, Fig. 4a, magenta dots).

**Figure 4.**
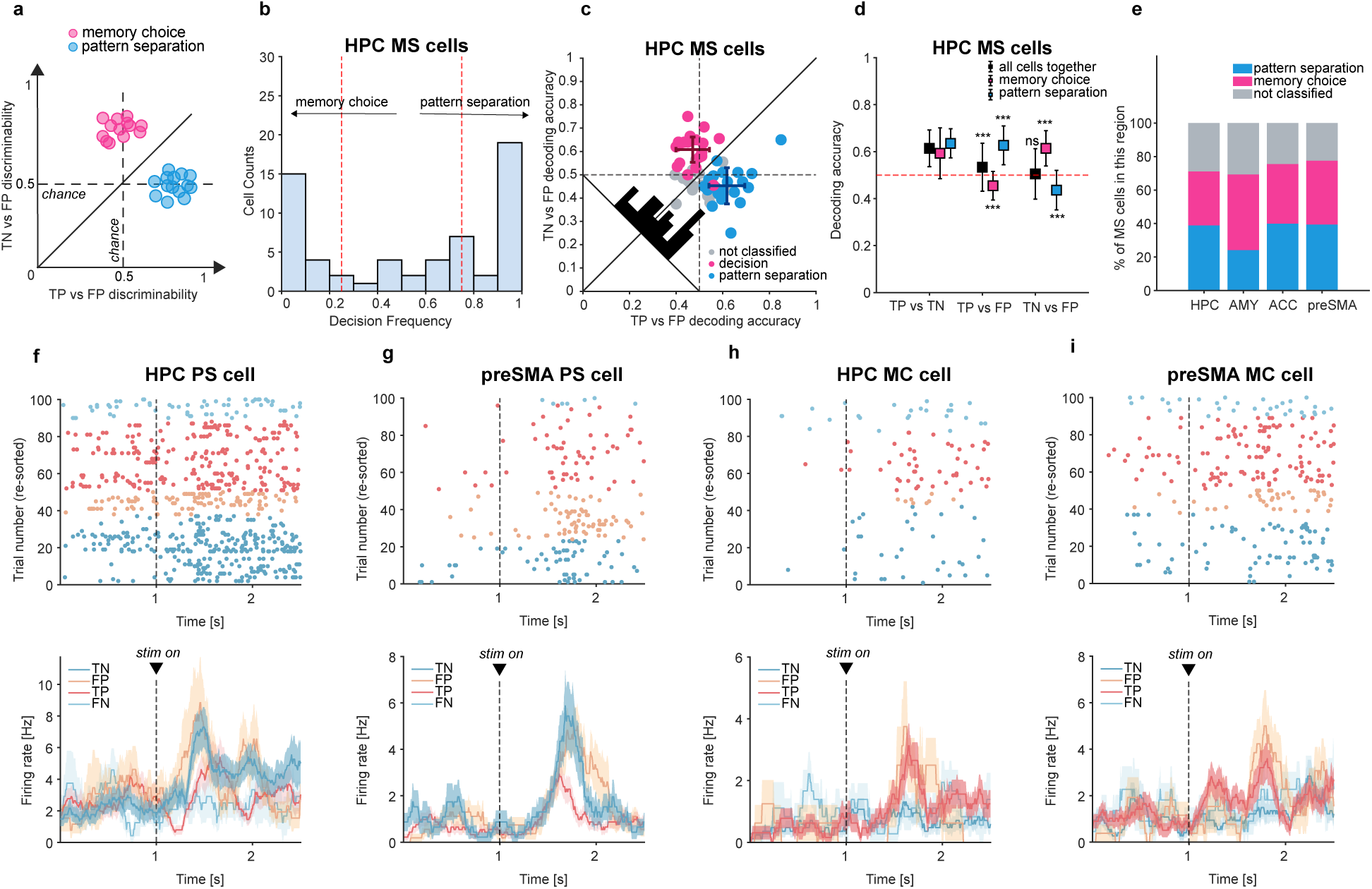
Activity of pattern separation subtype of MS cells distinguishes between true old and false old stimuli despite behavioral errors. Analysis of error trials. **a,** Hypothesis: Activity of MS neurons that reflect pattern separation can discriminate between true old (TP) and false old (FP) stimuli, despite the same behavioral choice being made, while they can not discriminate between true new (TN) and FP trials because they share the same ground truth (blue color). In contrast, neurons that signal choice differentiate stimuli with distinct decisions (TN and FP) but not stimuli that elicit the same decisions but different ground truth (TP and FP, pink color). **b,** Most MS cells in HPC (68.3%) are consistently defined as either the ‘memory choice’ or ‘pattern separation’ type, indicating the presence of two types of MS cells. Shown is the probability that a given MS cell is classified as one or the other across runs. Red line shows the criterion (75%) for a cell to be classified as one or the other subtype. **(c-e)** MS cells discriminate either between TP and FP or between TN and FP trials, but not both. **c,** Scatter plot of cross-validated single-cell decoding of TN vs FP trials (different choice, same ground truth) and TP vs FP (same choice, different ground truth). Each dot is a cell. Darker color is the mean ± s.d. for each decoding contrast. **d,** Same as c, but plotted as Mean ± sd. TP vs TN decoding is above chance by design since cells were pre-selected to distinguish correct new and old trials (i.e., MS cells). PS cells allowed above-chance decoding of TP vs FP (p=0.001, stimulus ground truth) while TN vs FP decoding was significantly below chance (p=0.001). MC cells allowed above-chance decoding of TN vs FP (p=0.001, behavioral choice) while TP vs FP decoding was significantly below chance (p=0.001). **e,** Proportion of PS and MC subtype of MS cells across brain areas. No significant difference in proportions was detected. **(f-i),** Example MS cells categorized as PS or MC subtype. Example ‘pattern separation’ (PS) cells in HPC (**f**) and preSMA (**g**). Example ‘memory choice’ (MC) cells in HPC (**h**) and preSMA (**i**). Note that PS cells, when they prefer novel stimuli (dark blue color), also respond to error trials that are new but were mistakenly recognized as ‘old’ (FP, beige color). In contrast, MC cells respond to trials with the same behavioral choice: ‘old’ correct in red and ‘old’ incorrect in beige

We used the same cross-validation procedure as described above to test this hypothesis. In each test/train split, the training trials were used for two purposes: to train a single-cell decoder to differentiate either TP from FP, or TN from FP, and to calculate a single-neuron ROC-AUC for the same contrast (see Methods). Each MS cell was classified as ‘memory choice’ (MC) if its AUC for TN vs. FP was higher than for TP vs. FP, or ‘pattern separation’ (PS) if its AUC for TP vs. FP was higher than for TN vs. FP (Fig. 4a). The decoding accuracy for both contrasts was then evaluated based on the held out test trials. For each MS cell, this procedure yielded two decoding performance measures and the proportion of runs where it was classified as an MC or PS cell (Fig. 4b, S4a-c).

Most of the MS cells were classified as either the MC or PS type (72.1% across four brain areas, Fig. 4b, S4a-c), indicating that there are two distinct types of MS cells. We then performed further analysis for all MS cells (based on left out trials) that were classified as either type in at least 75% of the runs (Fig. 4f-i shows examples). Only cells that were classified as PS allowed significant decoding of TP vs. FP in left-out trials (Fig. 4c, d blue, x axis; figure shows HPC: 0.62±0.08 (Mean±s.d), p=0.001; for other brain areas see Fig. S4d-i). The choice decoding accuracy for PS cells was significantly below chance (Fig. 4c,d, blue, y axis; 0.45±0.08, p=0.001). This below-chance performance further supports the interprettion that the activity of this group of cells is reflective of pattern separation because it means that PS cells consistently encode FP trials according to their ground-truth identity rather than the subject’s choice, resulting in systematically flipped labels during choice decoding. This result shows that the response of this type of MS cells preserves memory fidelity despite patients making an error, i.e. their response is reflective of neuronal pattern separation.

Conversely, MC cells allowed above-chance decoding of behavioral choice (Fig. 4c,d, TN vs. FP: 0.61±0.07, p=0.001) while TP vs FP decoding was significantly below chance (Fig. 4d, 0.47±0.05, p=0.001). We observed generally the same pattern in all other recording sites: only PS cells allowed for significant decoding of memory ground truth from FP errors, while only MC cells allowed for significant decoding of patient choice (Fig. S4d-i). There was no significant difference in the proportion of MS cells that qualified as the PS/MC type across brain areas (Fig. 4e).

### Population-level representations of memory ground-truth and behavioral choice

Above analysis identifies signatures of pattern separation at the single neuron level. Was pattern separation particularly salient in the HPC, where it was long-hypothesied to originate^6,7^ or is it present just as strongly in other brain regions^39^? The single-neuron analysis was not able to provide an answer because pattern separation was seen in all brain areas examined. We reasoned that this is due to the highly correlated nature of the memory and choice signals, which differ only during error trials. We therefore next used a method to disentangle the two statistically using demixed principal component analysis (dPCA; see Methods)^40^. We defined population-level coding dimensions that separately captured neural variance due to memory ground truth and behavioral choice signals (Fig. 5a-e and 5f-j show examples of HPC and preSMA). These two coding dimensions and the interaction between the two explain a significant amount of variance in population activity (19%, 19%, and 18% respectively in HPC, Fig. 5b; 10%, 13%, 12% in preSMA, Fig. 5g; 18%, 19%, 20% in AMY, not shown; 18%, 20%, 20% in ACC, not shown).

**Figure 5.**
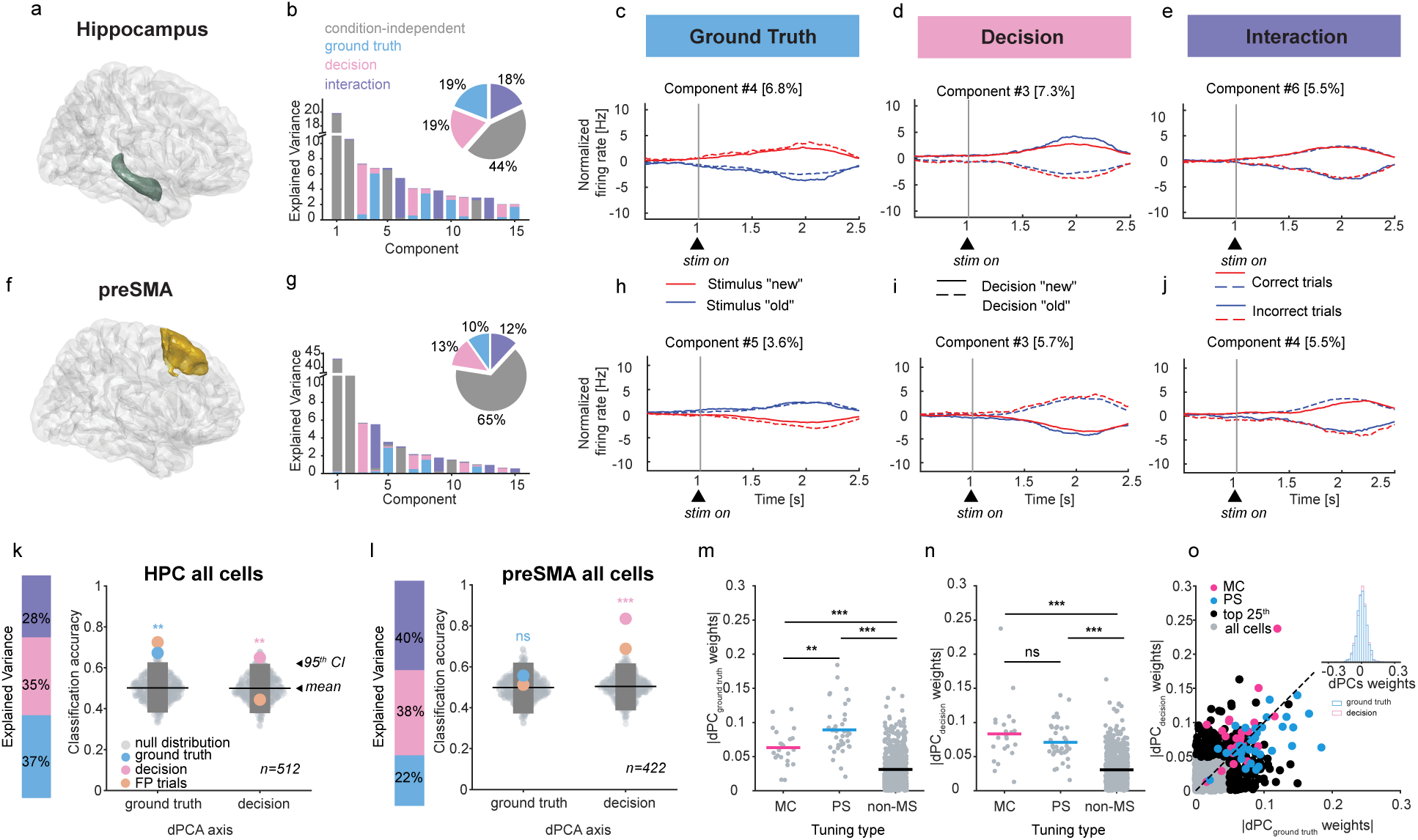
Both memory ground truth and behavioral choice is represented at the population level. **a-e,** Demixed principal component analysis (dPCA) of memory and choice representations in HPC. **a,** 3D brain model using the Brainnetome Atlas with highlighted HPC (green color). **b,** Both ground truth, decision, and the interaction between the two explain significant amounts of variance (19%, 19%, and 18%, respectively). **c-e,** Neural activity projected onto the ground truth (**c**), decision (**d**), and interaction (**e**) dPC explaining the most variance for each variable. Note the separability of memory ground truth (red vs blue colors) in c, behavioral choices (solid vs dashed lines) in d, and correct and incorrect trials in e. **f-j,** Same as a-e, but for preSMA. **k,** The bar on the left shows the variance explained by the dPCA parameters when only a single time window (1.5s after stim onset) is considered. In HPC, both memory ground truth and decision can be decoded at the single-trial level from the corresponding dPCA axes (mean during 1.5s after stimulus onset). Only cells with a minimum of 5 trials in each condition were considered (n=512). In HPC, decoding of both ground truth (p=0.003) and decision (p=0.009) was significant. The beige dot marks FP trials classification accuracy. **i,** Same as k, but for preSMA. In preSMA only decision was significantly decodable (p=1.7×10-8, ground truth p=0.17). **m-o,** Relationship between population (dPCA result) and single-cell tuning in HPC. MC and PS cells had higher dPCA weights than non-MS cells for both ground truth (MS p=5.9×10-9, PS p=1.6×1–36), and decision (MC p=2.2×10-20, PS p=9.2×10-21) axes. PS cells also had significantly higher weights for ground truth decoder axis than MC cells (p=0.005, decision axis MC vs PS p=0.2). All m and n p-values are two-sample t-test. **o,** Same as m and n but plotted against each other. The top right corner insert is the distribution of dPCA weights, for ground truth (light blue) and decision (pink) axes. Note centering on 0. * p<0.05, ** p<0.01, *** p<0.00 1

Can the dPCA-defined dimensions reliably convey single-trial memory and choice information to a downstream readout population? To test this question, we next performed single-trail decoding of the responses projected onto the dPCA axes, averaged in a single 1.5s long time window that starts 200 ms after stimulus onset. To assess statistical significance, we used single-trial decoding on a single left out trial of each type (correct and incorrect trials for both ‘new’ and ‘old’ stimuli) in each fold (see Methods). Only in HPC was decoding accuracy significantly larger than expected by chance for both memory ground truth and choice (Fig. 5k, left and right; mean accuracy is 0.67, p=0.003, and 0.65, p=0.009, respectively; chance is 0.5). When we analyzed the decoding accuracy exclusively for left out FP trials (accuracy reported above is the average accuracy across all trial types: TP, TN, FP, FN), we found that they can be classified with high accuracy in HPC but only along the ground truth dPCA axis (Fig. 5k, left, beige dots, accuracy=0.72 vs. Fig. 5k, right, beige dots, accuracy=0.45). In contrast, in the preSMA, only behavioral choice was decodable (Fig. 5l, right, 0.83, p=1.7×10^−8^, ground truth 0.56, p=0.17). Directly comparing decoding accruacy between the two brain areas (see Methods) shows that in the HPC, memory ground truth can be decoded significantly better than in the preSMA (Δ(HPC, preSMA)=0.1, p=0.004, Fig. S6d). We could not decode better than expected by chance from either dimension in the AMY (Fig. S6a, 0.49, p=0.4 and 0.59, p=0.08) and the ACC (Fig. S6b, 0.4, p=0.06 and 0.57, p=0.13). Together, this analysis shows that memory ground truth information at the single trial level was only reliably decodable in the HPC at the population level. In contrast, in preSMA, only the subject’s behavioral choice was reliably decodable at the population level.

Finally, we examined whether the PS cells identified previously contributed to population-level decoding. To do so, we compared the weights assigned by dPCA for either coding dimension for cells classified as PS, MC, or non-MS cells. First, MC and PS cells had higher dPCA weights than non-MS cells for both the ground truth (Fig. 5m,o; MS p=5.9×10^−9^, PS p=1.6×10^−36^, two-sample t-test) and the decision (Fig. 5n,o; MC p=2.2×10^−20^, PS p=9.2×10^−21^, two-sample t-test) axes. Second, PS cells were assigned significantly higher weights along the ground truth axis than MC cells (Fig. 5m, p=0.005; decision axis: Fig. 5n, MC vs. PS p=0.2). Together, these findings suggest that hippocampal PS cells primarily carry the memory ground truth signal we observed at the population level, enabling downstream neural populations to discriminate between novel stimuli and highly similar stored memories.

### Pattern separation cells drive the continuous memory strength signal

How do pattern separation and memory strength relate to each other? We investigated how the activity of PS cells is related to memory strength to address this question. We first repeated the dPCA-based population analysis after excluding PS cells. This abolished the ability to decode ground truth from HPC, while leaving choice decoding unaffected (Fig. 6a, mean decoding accuracy for ground truth was 0.5, p=0.5; accuracy for choice decoding was 0.61, p=0.03, respectively). This was the case regardless of whether dPCA had access to all recorded cells (analysis so far) or only to MS cells (ground truth and choice mean decoding accuracy is 0.56, p=0.17 and 0.63, p=0.01, respectively). Due to this, we next used only MS cells as a group to further investigate their properties. The variance captured by dPCA differed significantly between MS cells in the HPC and preSMA. In the HPC, the majority of the explained variance in the firing rate of MS cells was attributed to stimulus ground truth (53%, Fig. 6b, top). In contrast, in preSMA, ground truth only accounted for 21% of the variance. Instead, the choice component accounted for the largest fraction of neural variance of MS cells (58%, Fig. 6b, bottom). This suggests that, while individual neurons outside HPC still carry ground-truth information, the largest source of trial-to-trial variability in areas outside the HPC (here, the preSMA) is behavioral choice, whereas in the HPC it is ground truth. These findings highlight area-specific differences.

**Figure 6.**
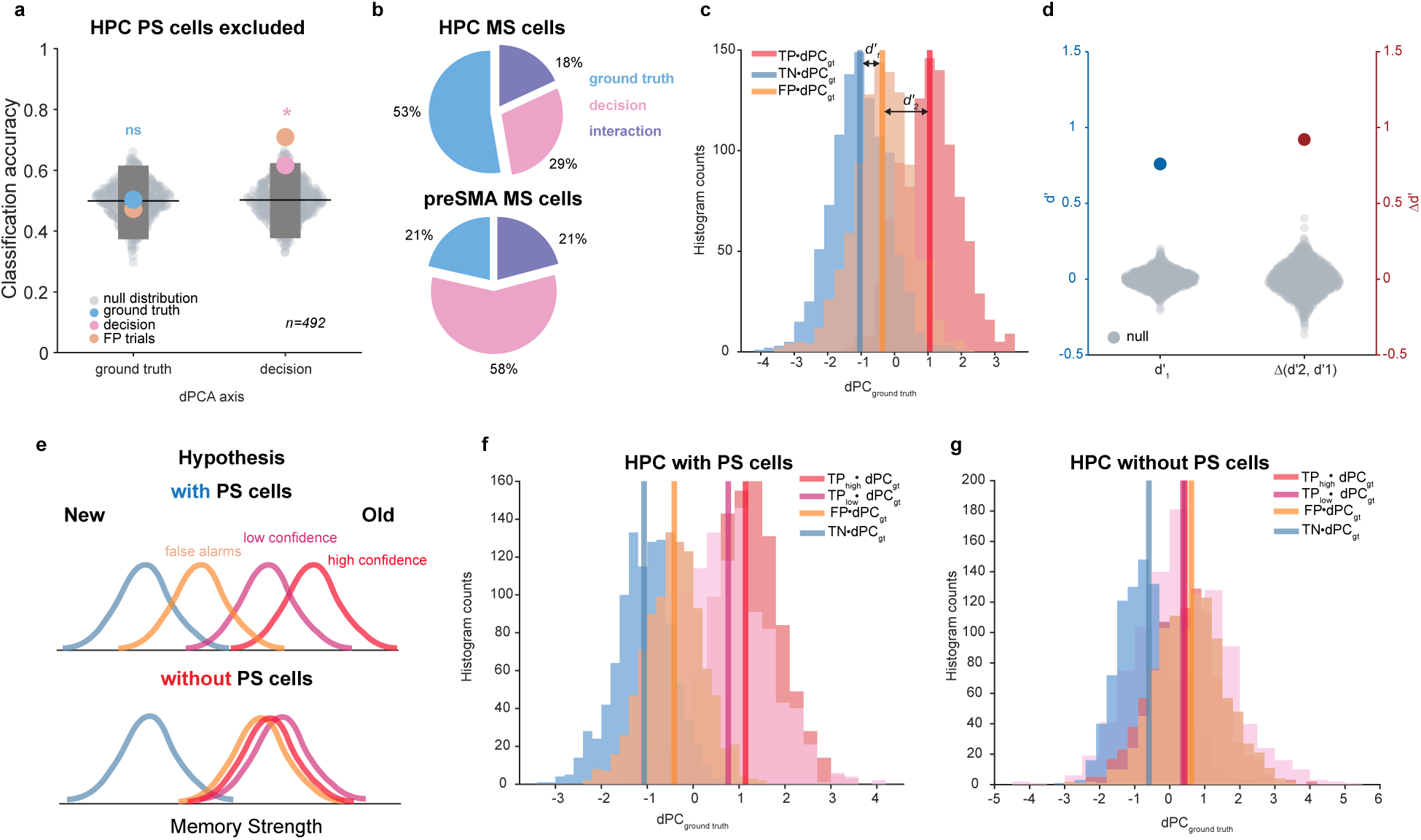
Pattern separation-related neural activity contributes to the continuous memory strength signal. **a,** Removing the PS-type MS cells (n=20) from the HPC population abolishes the ability to decode ground truth (p=0.5) but not decision (p=0.03). **b,** Majority of variance in population of all recorded MS cells is explained by ground truth (53%) and decision (58%) in the HPC and preSMA, respectively. Based on dPCA of all MS cells. **c,** Distribution of single-trial projections onto dPC ground truth decoder axis for TP, TN, and FP trials (red, blue, and beige colors). TP and FP distributions were more separable than FP and TN (d-prime = 1.68 (d’2) and 0.76 (d’1), respectively). **d,** Comparison of observed FP-TN d-prime (d’1) and difference in TP-FP (d’2) and FP-TN d-primes with the null distributions generated by calculating d-primes and taking the difference after shuffling trial labels. . None of the permuted values exceeded the observed values for both d’1 and d’2 - d’1 (both p=0.001, set as 1 divided by the number of permutations). **e,** Schematics of single detection theory (SDT) predictions. Top: memory strength represented as a continuous signal with recognition decisions based on whether this signal exceeds the decision threshold. On this axis, higher confidence decisions are associated with higher memory strength and would lie farther from the midpoint criterion. Bottom: without PS cells, no separation between ‘old’ decisions with different memory strength (high, low-confidence TP, FP) is achieved. **f,** Same as c, but with TP trials plotted separately for high and low confidence decisions. In HPC, the high-confidence TP (red color) distribution was the most separated from TN trials, with less separability for low-confidence TP and even less for FP trials, as STD would predict. When PS cells removed (**g**), projections of trials with different decisions are still separated, while high-confidence, low-confidence TP and FP trial projections (same decision) collapse on the ground truth axis. *** p<0.001

Why do patients still make mistakes, given that in their HPC, the response of neurons was different between FPs and TPs? We examined the distribution of projection values on the ground truth dPCA axis for individual trials to answer this question. We hypothesized that, despite being marked as ‘new’, FP trials are not identical to correct new (TN) trials and instead shifted more towards TP trials, potentially leading to weaker signals for downstream circuits to read out. To test this hypothesis, we constructed single-trial projection distributions by randomly leaving out one trial of each type and projecting it onto the ground truth dPCA axis estimated from the remaining trials (1000 iterations). We first computed d′ between FP and TN projection values (Fig. 6c, d′1) and confirmed that these distributions were separable (Fig. 6d, d′1=0.76, p=0.001 against shuffled label null). This result shows that the neural response during FP trials was not identical to that during TN trials. We then compared the separability of FP–TN and TP–FP (Fig. 6c, d′2) projection values and found that the TP–FP separation was larger than the FP–TN separation (Fig. 6d, d′2 = 1.68, Δ(d′2, d′1) = 0.92, p=0.001 against null distribution of differences). This response pattern indicates that FP trials remain closer to TN trials compared to TP trials but are nonetheless displaced toward TP along the dPCA ground truth axis. This result supports our hypothesis that FP trials, while being decoded as ‘new’, are less separable from TP trials along the memory ground truth axis compared to TN trials.

The placement of FP trials between TP and TN trials on the memory strength axis suggests that the ground truth signal in HPC could correspond to the signal detection theory-predicted memory strength signal. Within this framework, correct recognition decisions made with higher confidence are thought to lie farther away from lures than low-confidence decisions along the memory strength axis^41,42^. As PS cells enable memory ground truth decoding, we hypothesized that the activity of PS cells contributes to establishing the continuous memory strength signal in the HPC (Fig. 6e, top). If true, when PS cells are removed from the population, the different types of ‘old’ trials (FP, high and low confidence TP), each of which are accompanied by different memory strengths, should become indistinguishable from each other but still be differ from ‘new’ trials (Fig. 6e, bottom). To test this hypothesis, we employed the same single-trial projection analysis as described above, but now we constructed separate distributions for high- and low-confidence TP trials. When including all MS cells, the high-confidence TP distribution was the most separated from TN trials, with less separability for low-confidence TP and even less for FP trials (Fig. 6f). Conversely, in preSMA, all trial projections on the ground truth axis overlapped (Fig. S7a), highlighting HPC-specificity. In the HPC, after removing PS cells, ‘new’ and ‘old’ decision trials could still be distinguished, but high- and low-confidence TP and FP trials become indistinguishable (Fig. 6g). As an additional control, we repeated the same analysis but excluded the choice-signaling type of MS cells, MC cells. Without MC cells, the distribution placement resembled that of all MS cells together (Fig. S7b), suggesting that PS cells play a key role in establishing the continuous memory strength signal specifically, an insight that was only possible because we separated MS cell into types reflective of pattern separation (PS) and those reflective of the choice (MC). This result suggests that PS cells contribute to the hippocampal graded memory strength signal.

### Ground-truth memory signal is attenuated inside seizure onser zone and in the left hippocampus

It is known that neurons located within the seizure onset zone (SOZ) carry weaker memory-related signals than those outside of the SOZ^43^. We therefore next examined if memory ground truth decoding ability was different between hippocampal neurons recorded inside or outside the SOZ. Out of 3506 cells recorded across all brain areas, 27% were recorded from electrodes localized to the SOZ (n=1016; temporal lobe for HPC and AMY, frontal lobe for ACC and preSMA; Fig. S8a). The proportion of cells located within the SOZ was higher in the MTL compared to the MFC (43% and 40% in HPC and AMY vs 3% and 8% in ACC and preSMA, Fig. S8b), which is expected given the prominence of medial temporal lobe semiology in our patients. This was true for all cells (Fig. S8b) as well as when analysis was restricted to only MS cells (Fig. S8c). We further classified the temporal lobe epilepsy SOZs into SOZs that included the hippocampal electrodes or not and localized all hippocampal cells accordingly (Fig. S8d for all cells and Fig. S8e for MS cells, PS-type of MS cells is shown with darker shades). Single-trial decoding analysis revealed that only cells recorded outside of the SOZ allow for significant ground truth decodig (Fig. S8f, decoding accuracy=0.64, p=0.019), with decoding inside the SOZ not better than expected by chance (Fig. S8f, decoding accuracy = 0.52, p=0.39). Directly comparing decoding accruacy between the cell number-matched outside and inside SOZ populations (see Methods) shows that memory ground truth can be decoded significantly better outside of SOZ than inside (Δ(outSOZ, inSOZ)=0.12, p=0.004, Fig. S8g). This result highlights the specificity of HPC in driving memory ground truth signals and underscores the clinical relevance of our findings.

Next, we examined if laterality contributes to the memory ground truth signal. Out of 950 HPC cells, 413 were recorded from the left HPC and 537 from the right (Fig. S8h). The proportions of MS cells were similar between left and right HPC (36 and 40 MS cells respectively, Fig. S8i). PS cells were common in the right HPC (56% (19/34) vs 27% (7/26), X^2^ test of proportions p=0.025, Fig. S8i) suggesting a greater contribution of the right HPC to pattern separation of visual information. Correspondingly, we could significantly decode memory ground truth only from the right HPC (Fig. S8j, decoding accuracy = 0.66, p=0.01; left accuracy = 0.52, p=0.43). Directly comparing decoding accruacy between the cell number-matched right and left HPC populations (see Methods) shows that memory ground truth can be decoded significantly better from the right HPC than from the left (Δ(right, left)=0.14 , p=0.004, Fig. S8k). This result further supports a preferential role of the right HPC in representing memory ground truth for visual stimuli.

## Discussion

We identified signatures of mnemonic pattern separation in the HPC by contrasting neural responses between false and true positives trials. Our major contribution is the identification of PS cells in the HPC, whose response differs between FP and TP trials. We posit that the response of these PS cells is a reflection of successful pattern separation. Supporting the importance of the signal carried by PS cells, removing PS cells from the population led to the collapse of the memory strength gradient carried by hippocampal cells as a population. Our data identifies a cellular substrate for pattern separation, a computation long hypothesized to be a key component of the memory machinery^6,7,9,10,44^.

From the point of view of correct trials, PS cells respond identicially to the canonical MS cells that have been characterized before^23^. But when examining the two types of errors patients can make in this task, it becomes clear that there are two types of such cells: those that signal the ground truth regardless of the decision made, and those that signal the decision made regardless of the underlying memory signal. It was the response during error trials that allowed us to differentiate between these two distinct processes both at the single-neuron as well as at the population level. Examining error trials also allowed us to identify distinct contributions of different brain areas. MS cells were present in numbers exceeding those expected by chance in all four brain areas that we examined. But from the point of view of error trials, it was only the population of MS cells in the HPC that allowed for reliable decoding of ground truth.

Our findings were only possible because of the very large dataset that we have accumulated for this task across 97 patients (n= 3506 cells). MS cells are relatively rare to begin with, with typically ∼10% of HPC cells qualifying. Of these, only 60% qualify as PS cells, leading to an overall prevalance of <6%. FP trials, in turn, are also rare (by definition), requiring large numbers of patients to study them. Our finding that, when viewed at this scale, PS cells become identifable is an example for a neural process that becomes understandable only once large numbers of cells are available for a given brain area and task.

It has long been appreciated that when subjects forget having seen an item (a false negative), memory signals in the HPC are often still present^45,46^. The presence of a memory signal in this instance is compatible with successful pattern completion, even if at the behavioral level subjects were not able to access this information. In contrast, less is known about the contribution of the HPC to FPs, which, we posit, are key to identifying signatures of pattern separation. In our task, the probability of a FP judgment can be predicted with high reliability based on the similiarity of the test image with all previously seen images. This indicates that FPs are related to existing memory content rather than due to noise or guessing, making the amenable to study^34^. However, fMRI studies have generally found that neural signals in the HPC do not differ between TP and FP trials^47,48^, and as a result thereof FP trials have not so far been used as a vehicle to examine pattern separation. Here we now show that a specific subset of neurons in the HPC respond differentiatly to FP and TP trials, a difference that at the macroscopic level of fMRI appears to not be visible^49^.

Our findig that signatures of pattern separation are specifically visible only when examining the subset of new trials that resulted in FPs also puts into context earlier work reporting signatures of pattern separation. In the classic fMRI paradigm in which pattern separation was first reported, participants view sequences of objects with exact repetitions, similar but nonidentical lures, and novel objects intermixed^16^. In this experiment, pattern separation was operationalized as a differential response to similar (lure) images versus exact repeats, a pattern of response that was observed in the CA3/DG. However, the task imposed no memory demands, leaving it unknown whether the FP rate would have been higher during the lures compared to the novel images. Our data shows that this comparison is critical, because it is the only possible way to differentiate between novelty signals (which are well known to exist in the HPC) and signatures of pattern separation (Fig. S5a-d). Furthermore, we found that memory-and choice signals are intermixed, likely making it necessary to rely on single-neuron resolution to distinguish the two types of processes from each other.

Prior works show that features in deep-vision networks predict human similarity judgments well^50,51^, and that increased stimulus relatedness — whether perceptual similarity of pictures or semantic closeness of words — drives false recognition in healthy subjects^48,52–54^. By collecting human similarity judgments (Fig. S2j) and using pretrained DNNs we confirmed that similarity of a given new (lure) test image with all previously seen images was correlated positively with FP rate (Fig. 2). Demonstrating this relationship for our stimulus set was important because it validated that our paradigm reliably elicited similarity-driven memory errors (e.g. failures of pattern separation).

The activity of PS cells played a critical role in establishing the continuous memory strength gradient in the HPC^41,42,55^. Along this gradient, the weakest memory strength was associated with new items and the strongest with old items correctly received with high confidence (TP high). In between, in this order, were FPs and correctly received items with low confidence (TP low). This way, memory strength predicts declared confidence, a key aspect of episodic memory^23^. Removing PS cells, which make up only ∼6% in the HPC, leads to a collapse of FPs, TP high and TP low trials, thereby abolishing these critical differences. Removing the cells that predicted choice, on the other hand, left the gradient intact. Based on this result, we posit that it is specifically PS cells, and therefore pattern separation, that establishes the continous memory strength gradient. This interpretation would also explain why, on highly similar trials, patients made FP errors despite signals being separate in the hippocampus: one interpretation of this finding is that the threshold for declaring an item ‘old’ was lower than the average activity for FP trials (Fig. 6e).

Consistent with previous reports of memory disruption by epilepsy, particularly if the SOZ is in the right HPC^43,56^, the pattern separation signal was impaired in cells recorded from the patients with SOZ involving the HPC (Fig. S8f,g). Such disease-related disruption further supports the idea that this computation is implemented by hippocampal neurons rather than being inherited from upstream areas. Moreover, these findings highlight mnemonic pattern separation as a cognitive function that may be particularly sensitive to being disrupted by epilepsy with hippocampal involvement and establish a potential neural correlate of this impairment. This finding may have broader clinical relevance, as pattern separation signals could potentially serve as a biomarker of hippocampal involvement in epilepsy, including the specific contribution of the right HPC. Lastly, our finding that pattern separation is strongest in the right HPC is compatible with our task involving visual stimuli, indicating that measuring strength of pattern separation is a sensitive enough metric to reveal hemisphere-specific processes^57–62^. It remains an important future question whether repeating our experiment with verbal stimuli would reveal preferential involvement of the left hippocampus^63^.

An important open question is what computational process gives rise to the reponse of PS cells. Here, we posit that the activity of PS cells is a reflection of pattern separation between the current sensory input and previously seen images stored in memory. But PS cell activity is only the result of this process, rather than the process itself because they do not signal the content of the memory or sensory input as such. One candidate computational mechanism is offered by the HebbFF model^64^, in which stimulus representations remain fixed while memory is stored in synapses for which plasticity is triggered when encountering novel stimuli, providing a biologically plausible mechanism for generating familiarity signals. This stands in contrast to classical accounts of recognition memory and pattern separation, including Marr’s theory of memory computations and subsequent models of conjunctive coding in HPC^5–7,65^. These theories are concerned primarily with how sensory representations themselves are transformed (e.g. decorrelated) to facilitate storage and retrieval of memories, rather than with how a comparison between a current representation and previously stored representations is computed. Our findings therefore motivate computational models that operationalize memory via two separate representations: one for stimulus content and one for familiarity^64^. Within such a framework, PS cells may constitute a neuronal substrate for the latter, providing experimental evidence for a ground-truth memory signal that is dissociable from sensory representations themselves.

Another question that remains for future work is to examine whether mnmeonic pattern separation properties differ along the long axis of the hippocampus, as others have suggested^66,67^. Our recordings are predominantely from the anterior hippocampus, leaving open this question.

Finally, our analyses did not examine wheather pattern separation, including remapping, is present in stimulus content-selective representations in the MTL^20,68–72^. Existing literature suggests that concept cell representations are highly invariant, i.e. they do not change their activity in response to similar stimuli of the cell preferred concept^24^. Our experiment does not address this question because it was not designed to identify concept cells. Further studies should examine how familiarity signals interact with invariant stimulus representations during memory tasks with simulataneous recordings of concept and PS cells.

## Methods Participants

### Patients

A total of 97 patients with intractable epilepsy (133 sessions) participated in this study; of these, 59 (87) were re-analysis of previously published data^73^ and 38 (46) are newly recorded. Recordings from all 97 patients in the MFC are unpublished. We included only the sessions in which patients performed sufficiently well and in which they used both high and low confidence ratings for both new and old responses (Behavioral AUC >= 0.6). Based on these behavioral criteria, we excluded 13 patients (21 sessions), leaving 84 patients (112 sessions) for analysis (Supplementary Table 1).

The sample size was not predetermined using statistical methods, but the number of patients included is large compared to similar studies. All participants gave informed consent and volunteered to take part in the research^74^. The study protocols were approved by the institutional review boards of Cedars-Sinai Medical Center, Toronto Western Hospital, and Johns Hopkins Hospital.

### Online participants

Several independent pools of participants performed a multi-arrangement task^75^. First, 45 individuals were recruited through Prolific (www.prolific.com) to judge similarities between randomly selected 25 images that subsequently were used to evaluate model performance in explaining human perceptual similarity. Another group of participants judged similarities for images drawn from the same semantic category. In this version, each category subset contained 30 images within the category plus 4 scaffold images from different categories, which were later used to align distances across the entire dataset. For this, 16-20 participants were recruited for each category from task variant 3 (see below for task description): n = 20 for animals, n = 20 for food, n = 20 for cars, n = 20 for people, n = 16 for landscapes.

### Psychophysics

**Recognition memory task** (epilepsy patients). The task was the same as previously published^23,73,76^. It consists of two blocks—learning and retrieval—separated by a 10–30 min long delay. During learning, patients viewed 100 unique novel images. During retrieval, 50 of these images (old/familiar) were intermixed with 50 new images (novel). Patients judged each image as novel or familiar using a 1–6 confidence scale (Fig. 1a). Only data from the retrieval block are reported here. Prior to the main experiment, participants completed a short training version with a separate stimulus set. Three task variants were used, identical in design but employing different stimuli. For those patients who completed multiple sessions, recordings were made on different days using different task variants. All images subtended 9°×9°. Each trial consisted of image presentation, followed by a 0.5 s blank screen and then a response screen, which remained until an answer was given (Fig. 1a). We had 3 variants of this task that had the same structure and differed only in stimuli. Stimuli were natural scene photographs spanning five categories (Fig. S1a-c), with equal numbers from each. The task was implemented in MATLAB using Psychophysics Toolbox^77^.

**Similarity-judgments task** (online participants). To acquire ground-truth human perceptual similarity ratings, we recruited participants with self-reported normal or corrected-to-normal vision through Prolific (www.prolific.com). Participants performed a multi-arrangement task implemented on the web-based Meadows platform (http://meadows-research.com). The task design was as originally proposed^75^.

Participants were asked to judge the similarities of images by arranging them within the circular arena based on visual similarity (Fig. S2c). Participants were instructed to drag and drop each image to place similar images closer together and dissimilar ones further apart. Images could be moved repeatedly until the participant was satisfied with the arrangement. Clusters of images from the same semantic category often emerged naturally, and participants were encouraged to refine the placement by adjusting positions to capture more fine-grained differences within categories.

When satisfied with the initial arrangement, participants pressed the ‘Done’ button, and it constituted one arrangement trial. The first arrangement trial always included all images in the set. To estimate the initial RDM, the squared Euclidean distance was computed for each pair of arranged images. On the subsequent trials, participants were presented with a subset of images as determined by the current estimate of RDM— lift-the-weakest algorithm for adaptive design of item subsets, as originally proposed for the multi-arrangement task^75^. The final RDM for each participant was derived as the weighted average of distances from all arrangements in which an image pair appeared (Fig. S2i — averaged RDM across all participants).

### Analysis of behavior

#### Memory

Based on behavioral choice, all recognition trials were classified into TP (correct old), true TN (correct new), FN (incorrect old), and FP (incorrect new, also referred to throughout the manuscript as ‘false alarms’). Retrieval performance was quantified using ROC) analysis. Each ROC point (Fig. 1g) corresponds to the hit rate (y-axis) and false-alarm rate (x-axis) at a given confidence level, with the lower-left point reflecting the highest confidence (“old, confident”). Behavior AUC (Fig.1h) was calculated by integrating the area under the ROC curve. Throughout the manuscript, we included only sessions with patient-level AUC > 0.6 and with responses spanning both high- and low-confidence ratings for ‘new’ and ‘old’ judgments. For Fig. 3d analysis, we selected sessions where patients accurately discriminated between high and low confidence memories (minimal accuracy for high 70% and low 55%)^23^.

#### Models

To analyze the contribution of perceptual similarity to memory performance, we used several pre-trained DNNs (Fig. S2b). DINOv2 is a self-supervised vision transformer (ViT) model that extracts general-purpose visual features without training labels. CLIP is a natural language supervised ViT-based model, trained to match text and images. AlexNet and ResNet50 are deep convolutional neural networks (CNNs) trained to categorize images. The “pixel-wise” model is the correlation distance between pixels for a pair of images. The “categories” model was constructed as an RDM with binary values by assigning a dissimilarity of 0 to image pairs within the same semantic category and 1 to pairs from different categories.

#### Model performance in explaining similarity judgements

For a systematic evaluation of model performances in explaining human perceptual similarity, we collected similarity judgments for a dataset of 25 images (five images for each of five categories) (Fig. S2j). The stimuli dataset was processed with models to obtain model RDMs (Fig. S2k). For CNNs and ViTs, dissimilarity was defined as one minus the cosine similarity between feature vectors from the final model layer (for CNNs, the layer preceding the classification head). Pixel-wise dissimilarity was computed as one minus Pearson correlation between the pixel values of each image pair. After defining model RDMs, we compared each model RDM with each human subject’s similarity RDM with randomized tiebreaking Spearman’s ρa^35^.

#### Regression analysis

To test the possibility that memory false alarms at least partially result from the false interference that the new images were similar to previously studied images, for each new image shown to epilepsy patients during retrieval, we computed the probability of incorrectly recognizing this image as “old” across all patients. We then regressed these probabilities against the minimum cosine distance between model’s deep feature vectors for this image and all images presented during learning. To account for the possible contribution of an image-intrinsic sense of familiarity to false alarms, as one of the regressors, we used ResMem, the CNN trained to predict image memorability^38^. For direct comparison of the false alarm rate with human judgments, we collected additional similarity judgments for the entire image set of task variant 3 with independent participants judging different categories (also see ‘Participants’; Fig. S2m). As a regressor in this case, we used the minimum Euclidean distance between the new image and all same-category images presented during learning.

### Electrophysiology

Recordings were performed with hybrid (macro-micro) depth electrodes with embedded microwires (AdTech Medical)^78^. Implantation sites were determined based purely on clinical requirements and included various combinations of the HPC, AMY, ACC, and preSMA. Most patients received bilateral implants. Continuous broadband signal (0.1 Hz–9 kHz) was recorded from each microwire at a sampling rate of 32 kHz using the ATLAS system (Neuralynx) or 30 kHz (Blackrock Neurotech system) depending on the institution. All patients included in this study had at least one well-isolated single neuron in one or more of the targeted brain regions.

### Spike detection and sorting

Raw electrical signals were band-pass filtered in the 300 Hz–3 kHz range. Spike detection and sorting were performed with the template-matching semiautomated algorithm OSort v4.1^79^. All sorting results were manually inspected, and putative single units were identified and retained for further analyses. The quality of isolated units was quantitatively assessed using established metrics ^80^, including the proportion of inter-spike interval violations (<3 ms), waveform signal-to-noise ratio, projection distance between cluster pairs, and isolation distance of each cluster relative to all other detected spikes. Only neurons meeting these criteria were included in subsequent analyses. Additionally, we excluded the neurons with a firing rate less than 0.2 Hz, and those that were recorded from the electrodes localized outside of the target area (see below).

In total, we recorded from 4166 well isolated units across all brain areas (HPC n=1093, AMY n=1359, ACC n=826, and preSMA n=888). Based on behavioral performance, we excluded 660 units (see ‘Participants’ for exclusion criteria/session count) leaving a total of 3506 units for further analysis (950, 1187, 684, and 685 cells in HPC, AMY, ACC, preSMA, respectively).

### Electrode localization

Electrode localization was performed by co-registering postoperative computed tomography (CT) scans with preoperative magnetic resonance imaging (MRI) using FreeSurfer v5.3.0 and v7.4.1 as previously described^81^. Electrode coordinates were subsequently aligned to the MNI152 space and mapped to the CIT168 probabilistic atlas^82^ for standardized reporting and visualization using Advanced Normalization Tools v2.1. Final confirmation of electrode placement in gray matter was based on visual inspection of subject-specific MRI–CT co-registrations, rather than atlas-based visualization.

### SOZ localization

The SOZ location was determined based on the results of intracranial monitoring available for each patient at the end of they hospital stay (Supplementary Table 1). We localized each recorded neuron inside or outside SOZ based on whether a particular patient had an SOZ in that hemisphere of the MTL for HPC and AMY or MFC for ACC and preSMA. All neurons recorded in subjects with bilateral SOZs were determined to be inside the SOZ. This resulted in 406/950 and 476/1187 in HPC and AMY vs 18/684 and 55/685 in ACC and preSMA neurons to be localized inside SOZ. For comparison of ground truth decoding (see below) between inside and outside SOZ HPC, we localized all HPC cells inside or outside SOZ based on whether a particular patient had an SOZ that included HPC electrodes. This resulted in 396 cells localized outside the SOZ, 398 inside and 156 with unknown exact location (Fig. S8d). Similarly, 36 MS cells were recorded outside the SOZ, 26 inside and 14 unknow (Fig. S8e).

### Selection of neurons

Spikes were counted in a 1,5 s window starting 200 ms after stimulus onset. Memory-selective (MS) neurons were defined as those showing a significant difference between correctly identified novel and familiar stimuli within this window (p-value < 0.05, two-tailed bootstrap comparison of means, 1,000 iterations). MS neuron was classified as familiarity-selective (FS) if the mean firing rate for old trials exceeded that of new trials. If the neuron firing rate for new trials exceeded that of old, the MS cell was classified as novelty-selective (NS, relevant for Fig. 3d, e). Visually selective (VS) neurons were identified using a one-way ANOVA (factor: visual category, 5 levels) on the same spike counts (p-value < 0.05).

### Inclusion criteria for neurons

To be included in the analysis, we required a cell to have a mean firing rate of >0.2 Hz during the stimulus presentation period across all trials. Additionally, for each specific analyses, we included only cells that had a specific minimum number of trials of each possible type (typically 2 for ROC analysis and 5 for decoding analyses). The required minimum number of trials is specified in later sections describing each method.

FP rates are a function of stimulus category, with pictures that belong to some categories (such as ‘animals’) more frequently reported as FPs than for other categories (Fig. S1a-c for task variants 1-3). For this reason, the MS cells that in addition also code for visual category^20,23^ might be a confound for our analysis. While MS cells that are also category tuned are rare (1.9% (18/950) 1.85% ( 22/1187) 1.5% (10/684) 1.6% (11/685) in HPC, AMY, ACC, preSMA, respectively), we nevertheless excluded them for all analyses following Fig. 3g; Fig. 3c shows an example of such a cell) as a pre-caution.

### Chance level for cell selection

To estimate chance levels for cell selection, the procedure for identifying memory-selective cells was repeated after randomly permuting the trial labels (correct new/old) 500 times.

### Single-neuron analysis

To construct peristimulus time histograms (PSTHs, Fig. 3a,b; Fig. 4f-i) we count spikes in 200 ms sliding-window with 16 ms step size.

**Single-neuron ROC analysis.** Neuronal ROC curves were computed from spike counts in a 1.5 s window starting 200 ms after stimulus onset. The detection threshold was varied between the minimum and maximum observed spike counts, linearly spaced into 20 steps. The the AUC was obtained by integrating the area under the ROC curve.

We performed ROC analysis to quantify how well individual MS neurons can distinguish between different trials of interest (see below).

**High and low confidence comparison** (Fig. 3d, e). To assess how MS neurons differentiated novel and familiar trials across confidence levels, only neurons with at least two trials at each of the four confidence levels were analyzed. For fair comparison, one group in the ROC analysis was stratified by confidence, while the other was kept fixed. For FS neurons, all TN trials (regardless of confidence) served as the fixed group and were compared with high- and low-confidence TP trials. For NS neurons, all TP trials were fixed and compared with high- and low-confidence TN trials.

**Analysis of error trials** (S5b, also relevant for Fig. 4a-e). To assess the pattern of MS cell responses to trials with failed behavioral pattern separation (FP), we calculated AUC for pairs of trials with the same behavioral choice but different ground-truth (TP vs FP) or different choice but same ground truth (TN vs FP). Only neurons with at least 5 incorrect trials were included in this analysis.

**Single-cell decoding** (Fig. 3, 4). Single-cell decoding was performed using a Poisson naïve Bayes classifier. For each neuron, we used spike counts as features. The expected spike count (λ) was estimated as the mean across training trials for each class separately. Then, for each test observation, the log-likelihood was computed using Poisson distributions parameterized by the corresponding λ values, and class membership was assigned based on this likelihood. On each decoding run (n=100), we randomly left out one trial per class (2 trials total) for testing. The reported performance is the average across classifications of these 2×100 trials. To estimate the null distribution for decoding performance, we repeated the same decoding procedure but after shuffling the class labels 1000 times.

For group-level significance testing, we computed the null distribution for each cell, then averaged chance performances across cells for each permutation, resulting in a 1×1000 null distribution. P-value was set to the number of null values exceeding the mean decoding performance across cells. If no null values exceeded the observed performance, p-value was set to one divided by the number of permutations (1/1000).

**Error trials analysis** (Fig. 3g,h). To test if MS cells can discriminate FP from TP trials and compare this discriminability to the correct trials discriminability (TP vs. TN) in a fair way, we analyzed all recorded cells. To be included, the cell should have at least 5 FP trials. We count spikes in the 1.5s window starting 200 ms after stimulus onset. We used the same cross-validation procedure as described above. Training trials were used to train the decoder to differentiate either TP from TN, or TP from FP, and to select for MS cells as described above. For each neuron, this yielded two decoding performance measures and the proportion of runs where it was classified as an MS cell. We then compared the decoding performances for cells that were classified as MS in at least 75% of runs. Varying the MS cells classification threshold to 85 or 95% did not change the results reported in the main text (Fig. S3f,g).

**MS cell classification** (Fig. 4a-e). To assess if an MS cell carried a memory choice or memory ground truth signal, we analyzed MS cells with at least 5 FP trials. We count spikes in the 1.5s window starting 200 ms after stimulus onset. We used the same cross-validation procedure as described above. Training trials were used to train the decoder to differentiate either TP from FP (same choice, different ground truth), or TN from FP (different choice, same ground truth), and calculate AUC for the same pairs (also see **‘Single-neuron ROC analysis. Analysis of error trials’)**. Each MS cell was classified as MC if its AUC for TN vs. FP was higher than for TP vs. FP, or PS if its AUC for TP vs. FP was higher than for TN vs. FP. For each MS neuron, this yielded two decoding performance measures and the proportion of runs where it was classified as an MC or PS cell. We then compared the decoding performances for cells that were classified as either type in at least 75% of runs.

### Population analysis

#### Demixed principal component analysis (dPCA)

We used dPCA as described previously^40^. First, for each neuron, we computed the firing rate trace in a 500 ms moving window with a 16 ms step size and z-scored it with respect to the baseline (1 s time window before stimulus onset) mean and standard deviation. The dPCA algorithm begins by decomposing population activity into marginalized data matrices corresponding to the variables of interest. We constructed marginalized data matrices with respect to stimulus ground truth and behavioral choice. Note that correct trials (TP and TN) always have the same labels in both marginalizations, while incorrect (FP and FN) have opposite labels in ground truth versus choice. For each marginalized average, the algorithm next computes separate encoding and decoding matrices using regularized reduced-rank regression. The columns of the decoding matrix were used as the demixed principal components (dPCs), onto which N-dimensional data (single-trial data for testing and trial-averaged data for training) were projected. The values of this matrix quantify each neuron’s contribution to the marginalized representation.

For decoding analysis (Fig. 5k–o), we used data from a single time window (1.5 s starting 200 ms after stimulus onset), thereby removing variance related to time. Only neurons with an average firing rate of at least 0.2 Hz and with at least five trials per condition were included in the dPCA decoding analysis.

To assess the statistical significance of coding dimensions, we used a single-trial decoding procedure. For each iteration, one trial from each trial type (correct and incorrect trials for both ‘new’ and ‘old’ stimuli) was randomly selected to form a single-trial activity matrix. The remaining trials were averaged to construct the training data matrix, which was used to compute the dPCA coding dimensions. The held-out single-trial data were then projected onto the first coding dimension (capturing the largest variance in training data) and classified according to the nearest class mean. This procedure was repeated 1,000 times, and decoding accuracy was defined as the proportion of correct classifications across repetitions. A null distribution was generated by shuffling trial labels and repeating the decoding procedure 500 times. P-value was set to the number of null values exceeding the observed decoding performance.

As a control to ensure that decoding in HPC was not influenced by the larger number of available neurons (512 vs. 422; note that these numbers are lower than reported in Fig. 1e due to the exclusion of neurons with fewer than 5 trials of each type for this analysis), we randomly selected 75% of the HPC population (n = 384) and repeated the decoding 250 times. The results remained qualitatively similar, with significant decoding of both memory ground truth and choice in HPC (Fig. S6c; p = 0.009 and p = 0.018, respectively).

To directly compare two decoders (such as HPC and preSMA in Fig. S6d, inside vs outside SOZ and left vs right HPC in Fig. S8) we calculated the difference in decoding accuracy between them after neuron-count matching and compared this difference against the null distribution of decoding differences computed after trial label shuffling (n=500 shuffles). For that, on each decoder permutation (n=250) we randomly selected 80-85% of the smallest population (preSMA, n=340; in SOZ, n=170; left HPC, n=190) from both populations and repeated the same decoding procedure as above. At the same time, we shuffled the trial labels for 500 times and computed the null distribution of decoding performance differences. P-value was set to the number of null values exceeding the observed difference with a lower bound of 1/250 when no null value exceeded the observed value.

#### Single-trial projection

To estimate single-trial projection distribution along memory coding direction, we computed dPCs on pre-selected MS cells only in a single time window (1.5 s starting 200 ms after stimulus onset). We randomly held out one trial of each type as described above. We averaged the remaining trials and computed the dPCA coding dimensions on them. The decoder matrix of each dimension (the largest variance component for memory ground truth or choice) was then normalized to unit magnitude to estimate the coding vector. The held-out trials were projected onto this coding vector using the dot product. This procedure was repeated 1,000 times, producing a distribution of single-trial projections along each coding direction. The separation between distributions of different trial types was quantified using d-prime:

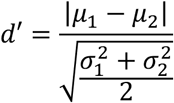

Where *μ*_1_ *and μ*_2_ are means of distributions 1 and 2, and *σ*_1_^2^ *and σ*_2_^2^ are variances of distributions 1 and 2.

We first tested if FP and TN distributions are separable (d′>0). For that, we computed d′ between FP and TN (Fig. 6c, d′1) and compared it against the null distribution of d-primes generated using the same procedure but after shuffling trial labels. On each leave-one-trial-out iteration (n=1000, see above) we shuffled trial labels for 500 times generating 1×500 single-trial projection distribution for each trial type and computed the shuffled d-prime. This resulted in 1000 null FP-TN d-primes against which we compared observed d′1. Since none of the null values exceeded the observed d-prime, the p-value was set to 1/1000 (dimensionality of null distribution).

We then compared the separability of FP–TN and TP–FP (Fig. 6c, d′2) distributions using the difference in corresponding d-primes. The positive difference between d′2 and d′1 would indicate that FP trials remain closer to TN than to TP but are nonetheless displaced toward TP along the dPCA ground truth axis. To test this statistically, we used the shuffled label distributions derived as described above and computed the difference in d-primes to generate the null distribution under the hypothesis that TP-FP distributions are not more separable than FP-TN distributions. This resulted in 1000 null d-prime differences. We then compared the observed difference in TP-FP d-prime and FP-TN d-prime with this null distribution. Since none of the null differences exceeded the observed difference in d-primes, the p-value was set to 1/1000 (dimensionality of null distribution).

For Fig. 6f the single-trial projection distributions were generated in a similar manner, but TP trials for testing (projecting) were selected based on their confidence (high or low). For Fig. 6g PS cells (defined as described in ‘**MS cell classification’** section) were excluded from the MS cell population.

## Acknowledgments

We are grateful to all members of the Rutishauser lab for valuable discussions, with particular thanks to Jake Gavenas. We thank Jasper van den Bosch from Meadows Research for guidance on similarity judgment collection. We thank the Cedars-Sinai, Toronto Western Hospital, and Johns Hopkins Hospital epilepsy monitoring unit physicians and staff for their support. We are deeply grateful the patients and their families for their time and participation. This project was funded by the NIH BRAIN Initiative (U01NS117839 to UR).

## Author contributions

N.K. and U.R. designed the study. N.K, C.M., J.D., Y.S., and U.R. collected the data. N.K. performed data analysis. S.C. localized electrodes. T.A.V, W.S.A, and A.N.M. managed patients and performed surgeries. C.M.R. provided patient care and supported data acquisition. N.K. and U.R. wrote the paper with input from all authors. U.R. acquired funding. U.R. and A.N.M. supervised the work.

## Data availability

MTL recordings of 59 patients are publicly available^73^. All MFC data and recordings from MTL and MFC from 38 patients (861 MTL and 1369 MFC neurons) is newly acquired and will be publicly released in the NWB format on DANDI upon acceptance of this manuscript.

## References

1. Standing, L. Learning 10000 pictures. Q. J. Exp. Psychol. 25, 207–222 (1973).

2. Brady, T. F., Konkle, T., Alvarez, G. A. & Oliva, A. Visual long-term memory has a massive storage capacity for object details. Proc. Natl. Acad. Sci. 105, 14325–14329 (2008).

3. Konkle, T., Brady, T. F., Alvarez, G. A. & Oliva, A. Scene Memory Is More Detailed Than You Think: The Role of Categories in Visual Long-Term Memory. Psychol. Sci. 21, 1551–1556 (2010).

4. Kramer, M. A., Hebart, M. N., Baker, C. I. & Bainbridge, W. A. The features underlying the memorability of objects. Sci. Adv. 9, eadd2981 (2023).

5. Marr, D. Simple memory: a theory for archicortex. Philos. Trans. R. Soc. Lond. B. Biol. Sci. 262, 23–81 (1971).

6. O’Reilly, R. C. & McClelland, J. L. Hippocampal conjunctive encoding, storage, and recall: Avoiding a trade-off. Hippocampus 4, 661–682 (1994).

7. Treves, A. & Rolls, E. T. Computational analysis of the role of the hippocampus in memory. Hippocampus 4, 374–391 (1994).

8. Rolls, E. T. Pattern separation, completion, and categorisation in the hippocampus and neocortex. Neurobiol. Learn. Mem. 129, 4–28 (2016).

9. Yassa, M. A. & Stark, C. E. L. Pattern separation in the hippocampus. Trends Neurosci. 34, 515–525 (2011).

10. Leal, S. L. & Yassa, M. A. Integrating new findings and examining clinical applications of pattern separation. Nat. Neurosci. 21, 163–173 (2018).

11. Quian Quiroga, R. No Pattern Separation in the Human Hippocampus. Trends Cogn. Sci. 24, 994–1007 (2020).

12. Suthana, N., Ekstrom, A. D., Yassa, M. A. & Stark, C. Pattern Separation in the Human Hippocampus: Response to Quiroga. Trends Cogn. Sci. 25, 423–424 (2021).

13. Quiroga, R. Q. How Are Memories Stored in the Human Hippocampus? Trends Cogn. Sci. 25, 425–426 (2021).

14. Rolls, E. T. On pattern separation in the primate, including human, hippocampus. Trends Cogn. Sci. 25, 920–922 (2021).

15. Quiroga, R. Q. Still challenging the pattern separation dogma: ‘quiero retruco’. Trends Cogn. Sci. 25, 923–924 (2021).

16. Bakker, A., Kirwan, C. B., Miller, M. & Stark, C. E. L. Pattern Separation in the Human Hippocampal CA3 and Dentate Gyrus. Science 319, 1640–1642 (2008).

17. Lacy, J. W., Yassa, M. A., Stark, S. M., Muftuler, L. T. & Stark, C. E. L. Distinct pattern separation related transfer functions in human CA3/dentate and CA1 revealed using high-resolution fMRI and variable mnemonic similarity. Learn. Mem. 18, 15–18 (2011).

18. Berron, D. et al. Strong Evidence for Pattern Separation in Human Dentate Gyrus. J. Neurosci. 36, 7569–7579 (2016).

19. Kumaran, D. & Maguire, E. A. Novelty signals: a window into hippocampal information processing. Trends Cogn. Sci. 13, 47–54 (2009).

20. Kreiman, G., Koch, C. & Fried, I. Category-specific visual responses of single neurons in the human medial temporal lobe. Nat. Neurosci. 3, 946–953 (2000).

21. Quiroga, R. Q., Reddy, L., Kreiman, G., Koch, C. & Fried, I. Invariant visual representation by single neurons in the human brain. Nature 435, 1102–1107 (2005).

22. Mormann, F. et al. A category-specific response to animals in the right human amygdala. Nat. Neurosci. 14, 1247–1249 (2011).

23. Rutishauser, U. et al. Representation of retrieval confidence by single neurons in the human medial temporal lobe. Nat. Neurosci. 18, 1041–1050 (2015).

24. Quiroga, R. Q. Concept cells: the building blocks of declarative memory functions. Nat. Rev. Neurosci. 13, 587–597 (2012).

25. Guzowski, J. F., Knierim, J. J. & Moser, E. I. Ensemble Dynamics of Hippocampal Regions CA3 and CA1. Neuron 44, 581–584 (2004).

26. Leutgeb, S., Leutgeb, J. K., Treves, A., Moser, M.-B. & Moser, E. I. Distinct Ensemble Codes in Hippocampal Areas CA3 and CA1. Science 305, 1295–1298 (2004).

27. Leutgeb, J. K., Leutgeb, S., Moser, M.-B. & Moser, E. I. Pattern Separation in the Dentate Gyrus and CA3 of the Hippocampus. Science 315, 961–966 (2007).

28. Lee, I., Yoganarasimha, D., Rao, G. & Knierim, J. J. Comparison of population coherence of place cells in hippocampal subfields CA1 and CA3. Nature 430, 456–459 (2004).

29. Lee, H., Wang, C., Deshmukh, S. S. & Knierim, J. J. Neural Population Evidence of Functional Heterogeneity along the CA3 Transverse Axis: Pattern Completion versus Pattern Separation. Neuron 87, 1093–1105 (2015).

30. Knierim, J. J. & Neunuebel, J. P. Tracking the flow of hippocampal computation: Pattern separation, pattern completion, and attractor dynamics. Neurobiol. Learn. Mem. 129, 38–49 (2016).

31. Neunuebel, J. P. & Knierim, J. J. CA3 Retrieves Coherent Representations from Degraded Input: Direct Evidence for CA3 Pattern Completion and Dentate Gyrus Pattern Separation. Neuron 81, 416–427 (2014).

32. Baker, S. et al. The Human Dentate Gyrus Plays a Necessary Role in Discriminating New Memories. Curr. Biol. 26, 2629–2634 (2016).

33. Mitchnick, K. A., Marlatte, H., Belchev, Z., Gao, F. & Rosenbaum, R. S. Differential contributions of the hippocampal dentate gyrus and CA1 subfield to mnemonic discrimination. Hippocampus 34, 278–283 (2024).

34. Parks, C. M., Yonelinas, A. P. & Wahlheim, C. N. Towards a unified theory of false memory for similar episodes. Psychon. Bull. Rev. 33, 88 (2026).

35. Schütt, H. H., Kipnis, A. D., Diedrichsen, J. & Kriegeskorte, N. Statistical inference on representational geometries. eLife 12, e82566 (2023).

36. Garoff-Eaton, R. J., Slotnick, S. D. & Schacter, D. L. Not All False Memories Are Created Equal: The Neural Basis of False Recognition. Cereb. Cortex 16, 1645–1652 (2006).

37. Norman, D. A. & Wickelgren, W. A. Strength theory of decision rules and latency in retrieval from short-term memory. J. Math. Psychol. 6, 192–208 (1969).

38. Needell, C. D. & Bainbridge, W. A. Embracing New Techniques in Deep Learning for Estimating Image Memorability. *Comput*. Brain Behav. 5, 168–184 (2022).

39. Amer, T. & Davachi, L. Extra-hippocampal contributions to pattern separation. eLife 12, e82250 (2023).

40. Kobak, D. et al. Demixed principal component analysis of neural population data. eLife 5, e10989 (2016).

41. Yonelinas, A. P. The Nature of Recollection and Familiarity: A Review of 30 Years of Research. J. Mem. Lang. 46, 441–517 (2002).

42. Wixted, J. T. Dual-process theory and signal-detection theory of recognition memory. Psychol. Rev. 114, 152–176 (2007).

43. Lee, S. J. et al. Single-neuron correlate of epilepsy-related cognitive deficits in visual recognition memory in right mesial temporal lobe. Epilepsia 62, 2082–2093 (2021).

44. Rugg, M. D. & Vilberg, K. L. Brain networks underlying episodic memory retrieval. Curr. Opin. Neurobiol. 23, 255–260 (2013).

45. Rutishauser, U., Schuman, E. M. & Mamelak, A. N. Activity of human hippocampal and amygdala neurons during retrieval of declarative memories. Proc. Natl. Acad. Sci. 105, 329–334 (2008).

46. Kirwan, C. B., Shrager, Y. & Squire, L. R. Medial temporal lobe activity can distinguish between old and new stimuli independently of overt behavioral choice. Proc. Natl. Acad. Sci. 106, 14617–14621 (2009).

47. Cabeza, R., Rao, S. M., Wagner, A. D., Mayer, A. R. & Schacter, D. L. Can medial temporal lobe regions distinguish true from false? An event-related functional MRI study of veridical and illusory recognition memory. Proc. Natl. Acad. Sci. 98, 4805–4810 (2001).

48. Slotnick, S. D. & Schacter, D. L. A sensory signature that distinguishes true from false memories. Nat. Neurosci. 7, 664–672 (2004).

49. Dubois, J., Berker, A. O. de & Tsao, D. Y. Single-Unit Recordings in the Macaque Face Patch System Reveal Limitations of fMRI MVPA. J. Neurosci. 35, 2791–2802 (2015).

50. Jozwik, K. M., Kriegeskorte, N., Storrs, K. R. & Mur, M. Deep Convolutional Neural Networks Outperform Feature-Based But Not Categorical Models in Explaining Object Similarity Judgments. Front. Psychol. 8, (2017).

51. Zhang, R., Isola, P., Efros, A. A., Shechtman, E. & Wang, O. The Unreasonable Effectiveness of Deep Features as a Perceptual Metric. Preprint at 10.48550/arXiv.1801.03924 (2018).

52. Roediger, H. L. & McDermott, K. B. Creating false memories: Remembering words not presented in lists. J. Exp. Psychol. Learn. Mem. Cogn. 21, 803–814 (1995).

53. Koutstaal, W., Schacter, D. L., Verfaellie, M., Brenner, C. & Jackson, E. M. Perceptually Based False Recognition of Novel Objects in Amnesia: Effects of Category Size and Similarity to Category Prototypes. Cogn. Neuropsychol. 16, 317–341 (1999).

54. Kim, J. & Yassa, M. A. Assessing recollection and familiarity of similar lures in a behavioral pattern separation task. Hippocampus 23, 287–294 (2013).

55. Squire, L. R., Wixted, J. T. & Clark, R. E. Recognition memory and the medial temporal lobe: a new perspective. Nat. Rev. Neurosci. 8, 872–883 (2007).

56. Reed, C. M. et al. Extent of Single-Neuron Activity Modulation by Hippocampal Interictal Discharges Predicts Declarative Memory Disruption in Humans. J. Neurosci. Off. J. Soc. Neurosci. 40, 682–693 (2020).

57. Milner, B. Psychological defects produced by temporal lobe excision. Res. Publ. Assoc. Res. Nerv. Ment. Dis. 36, 244–257 (1958).

58. Delaney, R. C., Rosen, A. J., Mattson, R. H. & Novelly, R. A. Memory Function in Focal Epilepsy: A Comparison of Non-Surgical, Unilateral Temporal Lobe and Frontal Lobe Samples. Cortex 16, 103–117 (1980).

59. Helmstaedter, C., Pohl, C., Hufnagel, A. & Elger, C. E. Visual Learning Deficits in Nonresected Patients with Right Temporal Lobe Epilepsy. Cortex 27, 547–555 (1991).

60. Spiers, H. J. et al. Unilateral temporal lobectomy patients show lateralized topographical and episodic memory deficits in a virtual town. Brain 124, 2476–2489 (2001).

61. Wilde, N. et al. WMS–III performance in patients with temporal lobe epilepsy: Group differences and individual classification. J. Int. Neuropsychol. Soc. 7, 881–891 (2001).

62. Burgess, N., Maguire, E. A. & O’Keefe, J. The Human Hippocampus and Spatial and Episodic Memory. Neuron 35, 625–641 (2002).

63. Camarillo-Rodriguez, L. et al. Temporal lobe interictal spikes disrupt encoding and retrieval of verbal memory: A subregion analysis. Epilepsia 63, 2325–2337 (2022).

64. Tyulmankov, D., Yang, G. R. & Abbott, L. F. Meta-learning synaptic plasticity and memory addressing for continual familiarity detection. Neuron 110, 544–557.e8 (2022).

65. Rolls, E. T. Functions of Neuronal Networks in the Hippocampus and Neocortex in Memory. in Neural Models of Plasticity 240–265 (Elsevier, 1989). doi:10.1016/B978-0-12-148955-7.50017-5.

66. Freelin, A., Wolfe, C. & Lega, B. Models of human hippocampal specialization: a look at the electrophysiological evidence. Trends Cogn. Sci. S1364661324003188 (2024) doi:10.1016/j.tics.2024.11.009.

67. To, T. V., Wang, D. X., Wolfe, C. B. & Lega, B. C. Neurophysiological evidence of human hippocampal longitudinal differentiation in associative memory. Nat. Commun. 16, 6845 (2025).

68. Suthana, N. A. et al. Specific responses of human hippocampal neurons are associated with better memory. Proc. Natl. Acad. Sci. 112, 10503–10508 (2015).

69. Kyle, C. T., Stokes, J. D., Lieberman, J. S., Hassan, A. S. & Ekstrom, A. D. Successful retrieval of competing spatial environments in humans involves hippocampal pattern separation mechanisms. eLife 4, e10499 (2015).

70. Dimsdale-Zucker, H. R., Ritchey, M., Ekstrom, A. D., Yonelinas, A. P. & Ranganath, C. CA1 and CA3 differentially support spontaneous retrieval of episodic contexts within human hippocampal subfields. Nat. Commun. 9, 294 (2018).

71. Reagh, Z. M. & Ranganath, C. Flexible reuse of cortico-hippocampal representations during encoding and recall of naturalistic events. Nat. Commun. 14, 1279 (2023).

72. To, T. V., Wang, D. X., Wolfe, C. B. & Lega, B. C. Neurophysiological evidence of human hippocampal longitudinal differentiation in associative memory. Nat. Commun. 16, 6845 (2025).

73. Chandravadia, N. et al. A NWB-based dataset and processing pipeline of human single-neuron activity during a declarative memory task. Sci. Data 7, 78 (2020).

74. Feinsinger, A. et al. Ethical commitments, principles, and practices guiding intracranial neuroscientific research in humans. Neuron 110, 188–194 (2022).

75. Kriegeskorte, N. & Mur, M. Inverse MDS: Inferring Dissimilarity Structure from Multiple Item Arrangements. Front. Psychol. 3, (2012).

76. Faraut, M. C. M. et al. Dataset of human medial temporal lobe single neuron activity during declarative memory encoding and recognition. Sci. Data 5, 180010 (2018).

77. Brainard, D. H. The Psychophysics Toolbox. Spat. Vis. 10, 433–436 (1997).

78. Minxha, J., Mamelak, A. N. & Rutishauser, U. Surgical and Electrophysiological Techniques for Single-Neuron Recordings in Human Epilepsy Patients. in Extracellular Recording Approaches (ed. Sillitoe, R. V.) 267–293 (Springer, New York, NY, 2018). doi:10.1007/978-1-4939-7549-5_14.

79. Rutishauser, U., Schuman, E. M. & Mamelak, A. N. Online detection and sorting of extracellularly recorded action potentials in human medial temporal lobe recordings, in vivo. J. Neurosci. Methods 154, 204–224 (2006).

80. Hill, D. N., Mehta, S. B. & Kleinfeld, D. Quality Metrics to Accompany Spike Sorting of Extracellular Signals. J. Neurosci. 31, 8699–8705 (2011).

81. Minxha, J., Adolphs, R., Fusi, S., Mamelak, A. N. & Rutishauser, U. Flexible recruitment of memory-based choice representations by the human medial frontal cortex. Science 368, eaba3313 (2020).

82. Tyszka, J. M. & Pauli, W. M. In vivo delineation of subdivisions of the human amygdaloid complex in a high-resolution group template. Hum. Brain Mapp. 37, 3979–3998 (2016).

