## Supplementary Material for "Cellular code for mnemonic pattern separation in the human hippocampus is revealed by false memories"

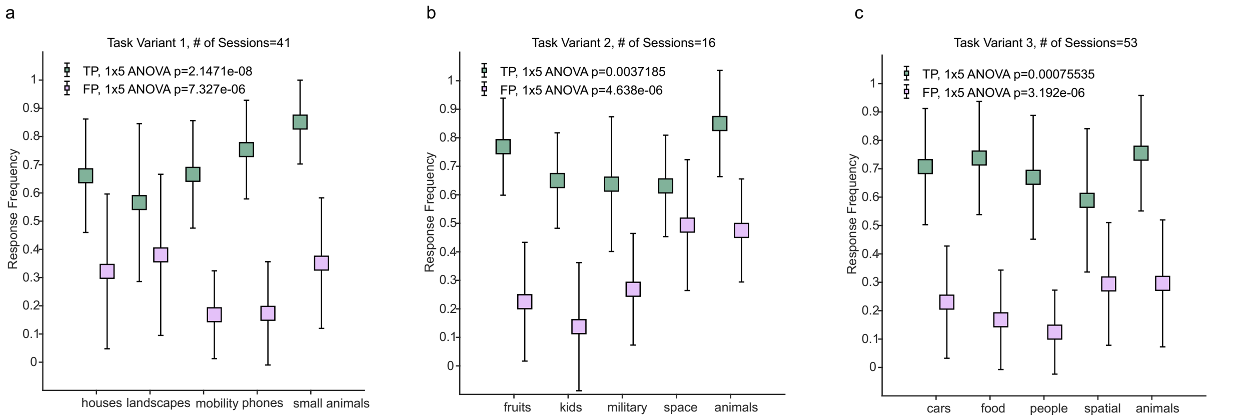


**Supplementary Figure 1. Additional behavior analysis.**
**(a-c),** Category-specific biases in correct and incorrect recognition responses. **a**, In task variant 1, different categories had significantly different proportions of true positive (TP, green color, 1x5 ANOVA p=2.1x10-8) and false positive (FP, light purple, 1x5 ANOVA p=7.3x10-6) responses. **b**, Same as (a), but for task variant 2. TP p=0.004, FP p=4.6x10-6. **c**, Same as (b), but for task variant 3. TP p=0.0008, FP p=3.2x10-6. Error bars are means across sessions ±s.d.

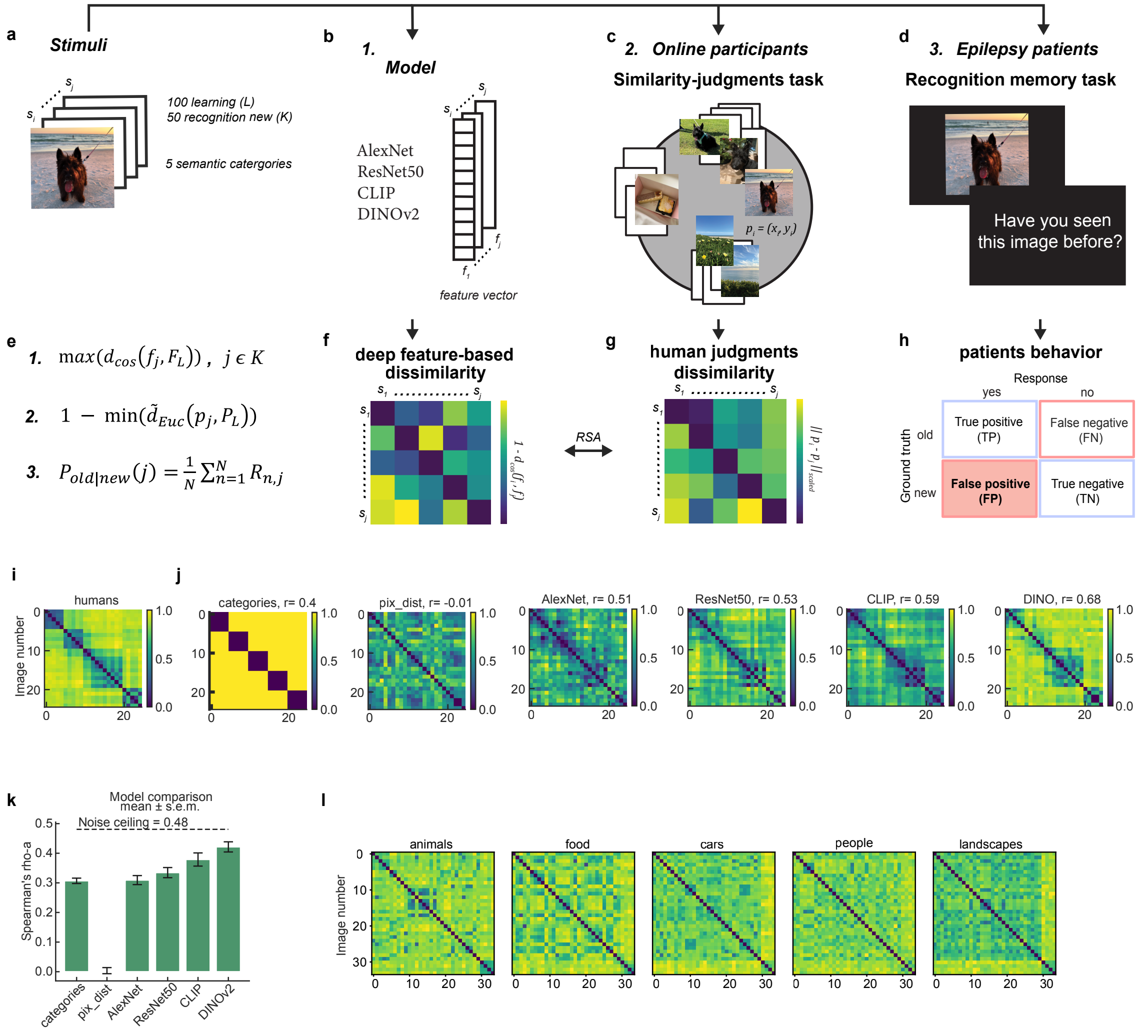

**Supplementary Figure 2. Visual and semantic deep neural networks reliably predict human perceptual similarity and memory false alarms.**

**(a-h)** Methods to measure similarity and false positives. **a,** Each visual stimulus was characterized by how similar it was to all other visual stimuli. **b,** Each image was parameterized with different pre-trained deep neural networks (DNNs, listed here) to extract feature vectors (*f*) from the last network layer. **c,** Multi-arrangement task to collect human similarity judgments for a subset of images (task variant 3). *p* denotes the 2D coordinates that each image *i* has on each trial after a participant placed it. **d,** Same images were used in a recognition memory task in a clinical setting. Each patient (n = 59) typically performed one task variant, while some did two variants or all 3 (n = 22 and n = 3). **e,** Metrics that were calculated from each experiment (1, DNN, 2, human similarity judgments, 3, recognition memory task) to evaluate the perceived similarity effect on memory false alarms (FP). **f,** Schematic DNN representational dissimilarity matrix (RDM) used in representational similarity analysis (RSA). **g,** Same as f, but for human similarity judgments from c. **h,** Confusion matrix of possible stimulus ground truth-patient response combinations; false positive (FP) trials used in the analysis presented on a red background. **i,** RDM of human judgements from **c**, averaged across all participants (n=45). **j,** RDMs of models used to evaluate human similarity judgments from c. *r* in the title of each plot is the Pearson correlation coefficient from correlation of model RDM with averaged humans RDM **(i)**. **k,** Model evaluation results in explaining human perceptual similarity. All models except for the pixel-wise model performed significantly better than 0 (one-sided t-test p-value<0.001, Bonferroni-corrected α=0.0013). CLIP, and DINOv2 significantly outperformed the category model with DINOv2 being significantly better than all other models (FDR-corrected pairwise t-test for 28 model-pairs comparisons). Only DINOv2 performance did not significantly differ from the lower bound noise ceiling (one-sided t-test p-value=0.0014). **l,** RDMs of within-category human similarity judgements (n=15 online participants per category). Each stimulus set contained 30 images from the same semantic category, along with 4 scaffold images (positions 31–34 in the RDMs) from different categories, which were later used to align distances across the entire dataset.

*dcos* cosine similarity, *fj* DNN feature vector of new recognition image *j*, *FL* feature vectors of all learning images L, *dEuc* min-max normalized Euclidean distance, *pj* multi-arrangements coordinates of new recognition image j, *PL* multi-arrangements coordinates of all learning images L, *Pold|new* probability of incorrectly recognizing new recognition image j as "old", *N* number of epilepsy patients performed the task (n = 41, 16, 53 for variants 1,2,3, respectively), *Rnj* response of patient n to image *j*.

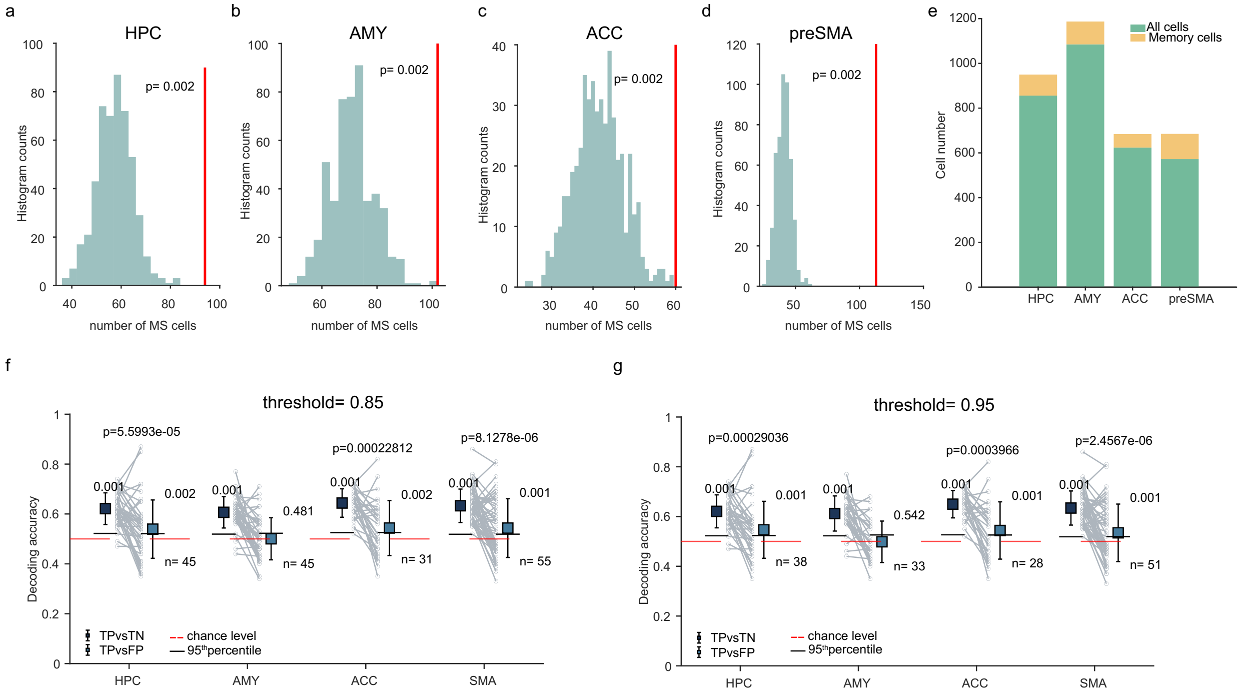


**Supplementary Figure 3. Additional analysis of memory selective (MS) cells.**
**(a-d)** Bootstrap statistics for the number of selected cells. In all brain areas (HPC (a), AMY (b), ACC (c), preSMA (d)) number of selected MS cells (red line) is larger than estimated chance values (grey). Null distribution is estimated by shuffling trial labels and re-running the selection procedure 500 times. The p-value was set to 1/500 (the number of permutations), because none of the estimated chance values exceeded the observed value. **e,** Number of isolated single units across recording sites (green, n total across 4 brain areas = 3137, excluding MS) and number of units classified as MS (yellow, n=369). **(f-g)**, Cross-validated single-cell decoding performance for neurons that were memory-selective in at least 85% (f) or 95% (g) of randomly selected trials. Same as Fig.3 h but using different thresholds to classify a cell as MS. Varying thresholds did not affect the conclusion that in all recording sites except for AMY, both TP vs TN and TP vs FP were decodable significantly above chance, with TP vs TN decoding being significantly higher than TP vs FP. Only the number of selected cells decreased as expected.

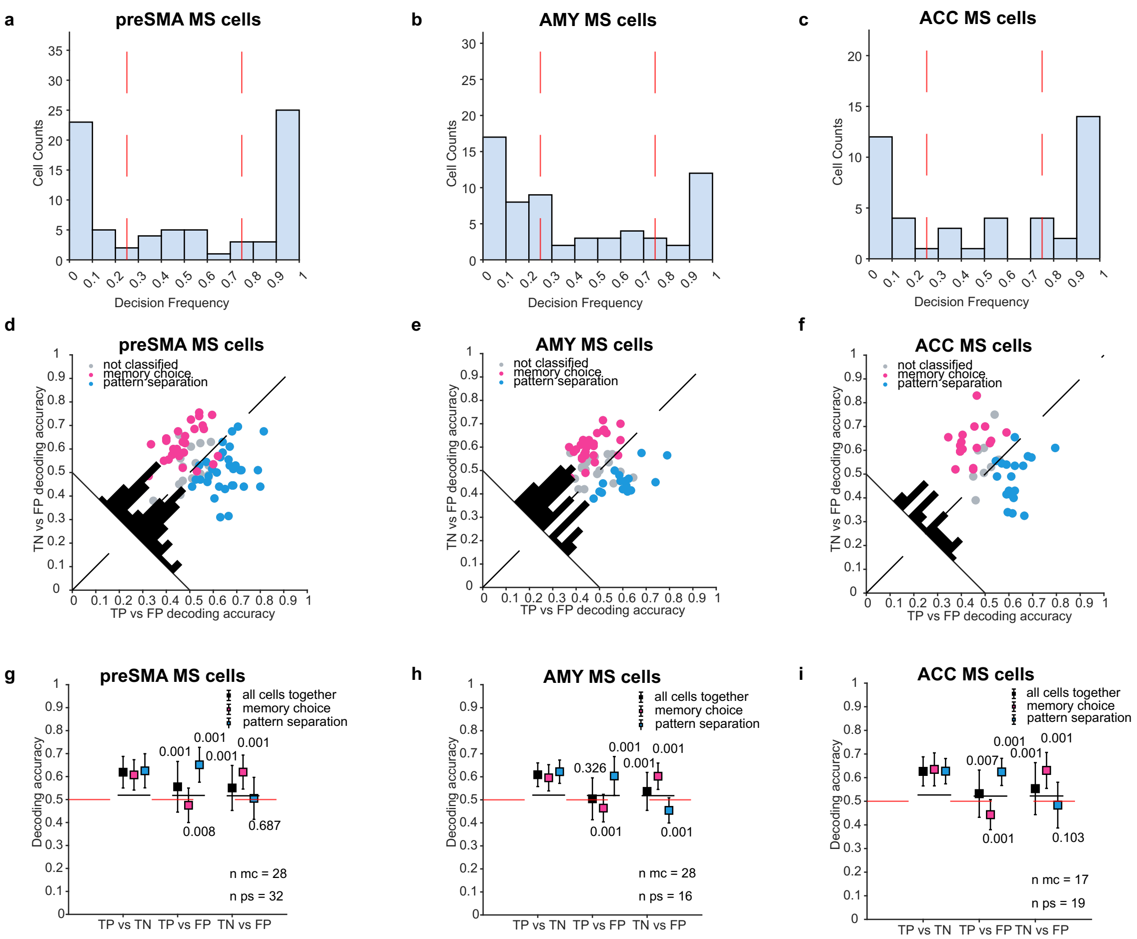


**Supplementary Figure 4. Analysis of error trials in other recording sites.**
**(a-c)**, Same as Fig.4 b but for preSMA, AMY and ACC. Most MS cells in preSMA (81.6%), AMY (63.5%) and ACC (73.3%) are consistently defined as either the 'memory choice' or 'pattern separation' type, indicating the presence of two types of MS cells. Shown is the probability that a given MS cell is classified as one or the other across runs. Red line shows criteria (75%) for a cell to be classified as one or the other subtype. **(d-f)** Same as Fig.4 c but for preSMA, AMY and ACC. Scatter plot of cross-validated single-cell decoding of TN vs FP trials (different choice, same ground truth) and TP vs FP (same choice, different ground truth). Each dot is a cell. **(g-i),** Same as (d-f), but plotted as Mean±sd. TP vs TN decoding is above chance by design since cells were pre-selected to distinguish correct new and old trials (i.e., MS cells). PS cells allowed above-chance decoding of TP vs FP ($p<0.01$, stimulus ground truth). MC cells allowed above-chance decoding of TN vs FP ($p<0.01$, behavioral choice) while TP vs FP decoding was significantly below chance ($p<0.01$) in all brain areas.

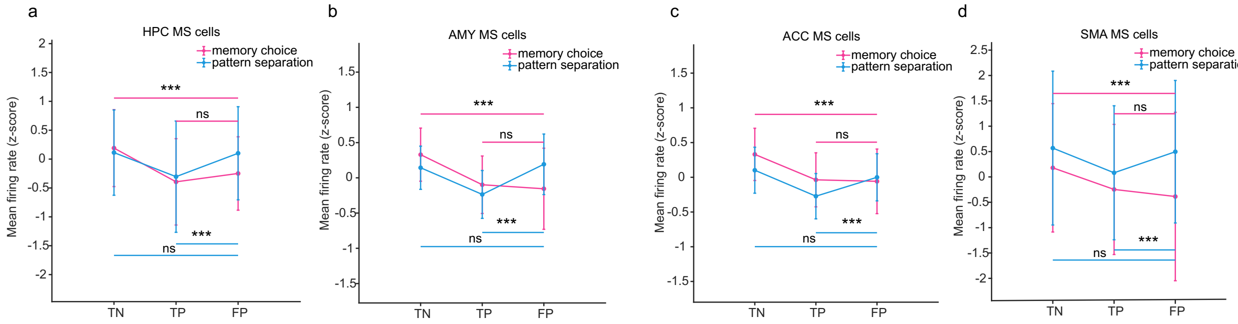


**Supplementary Figure 5. Comparison with methods to quantify pattern separation used in prior studies.**
**(a-d),** Comparison of baseline-normalized averaged firing rate for TN, TP, and FP trials for PS and MC cells. **a,** In HPC PS cells (blue) had significantly different firing rate for FP and TP trials (p=8.6x10-7) but not for FP and TN trials (p=0.7)—corresponding to previously proposed pattern separation-like response. MC cells (magenta) had significantly different firing rate for FP and TN trials (p=6.7x10-5) but not for FP and TP trials (p=0.4). **b,** Same as **a**, but for AMY: PS cells (blue) had significantly different firing rate for FP and TP trials (p=1x10-5) but not for FP and TN trials (p=0.7). MC cells (magenta) had significantly different firing rate for FP and TN trials (p=3.7x10-8) but not for FP and TP trials (p=0.8). **c,** Same as **a,** but for ACC: PS cells (blue) had significantly different firing rate for FP and TP trials (p=0.0001) but not for FP and TN trials (p=0.1). MC cells (magenta) had significantly different firing rate for FP and TN trials (p=0.0002) but not for FP and TP trials (p=0.5). **d,** Same as **a,** but for preSMA: PS cells (blue) had significantly different firing rate for FP and TP trials (p=3.2x10-8) but not for FP and TN trials (p=0.3). MC cells (magenta) had significantly different firing rate for FP and TN trials (p=1.5x10-8) but not for FP and TP trials (p=0.2). All p-values are Wilcoxon signed rank test. * p<0.05, ** p<0.01, *** p<0.001.

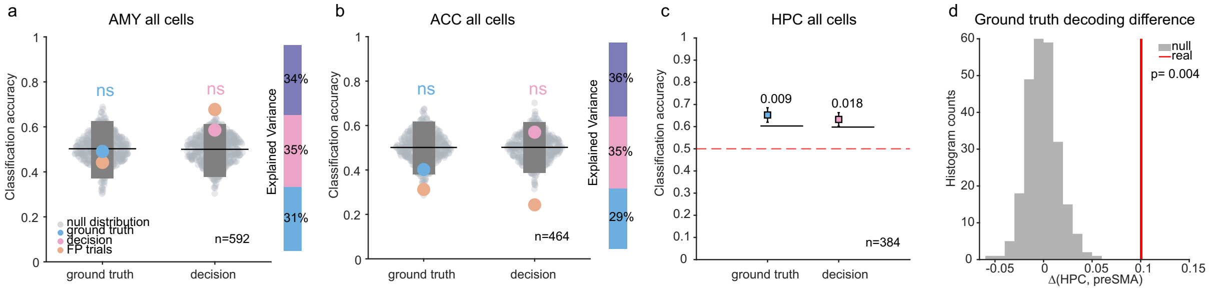


**Supplementary Figure 6. Additional dPCA population decoding analysis.**
In both AMY (**a**) and ACC (**b**), neither memory ground truth (p=0.4 and p=0.06 below chance) nor decision (p=0.08 and p=0.13) could be significantly decoded at the single-trial level from the corresponding dPCA axes (mean during 1.5s after stimulus onset). Only cells with a minimum of 5 trials in each condition were considered (n=592 AMY and n=464 ACC). The beige dot marks FP trials classification accuracy. The bar on the right shows the variance explained by the dPCA parameters when only a single time window (1.5s after stim onset) is considered. **c,** In HPC, both memory ground truth (p=0.009) and decision (p=0.018) can be decoded at the single-trial level from the corresponding dPCA axes even after downsampling to 384 cells (75% of HPC population). Error bars are means across runs (n=250)± s.d. Black lines are the 95th percentile of the null distribution. **d,** Ground-truth decoding in HPC is significantly higher than in preSMA (mean difference across n=250 subsamples, delta=0.1, red line). The null distribution (grey) is generated by shuffling the trial labels and repeating the same decoding procedure for both areas and taking the difference. None of the permuted values exceeded the observed difference (p=0.004, p-value set as 1 divided by the number of permutations).

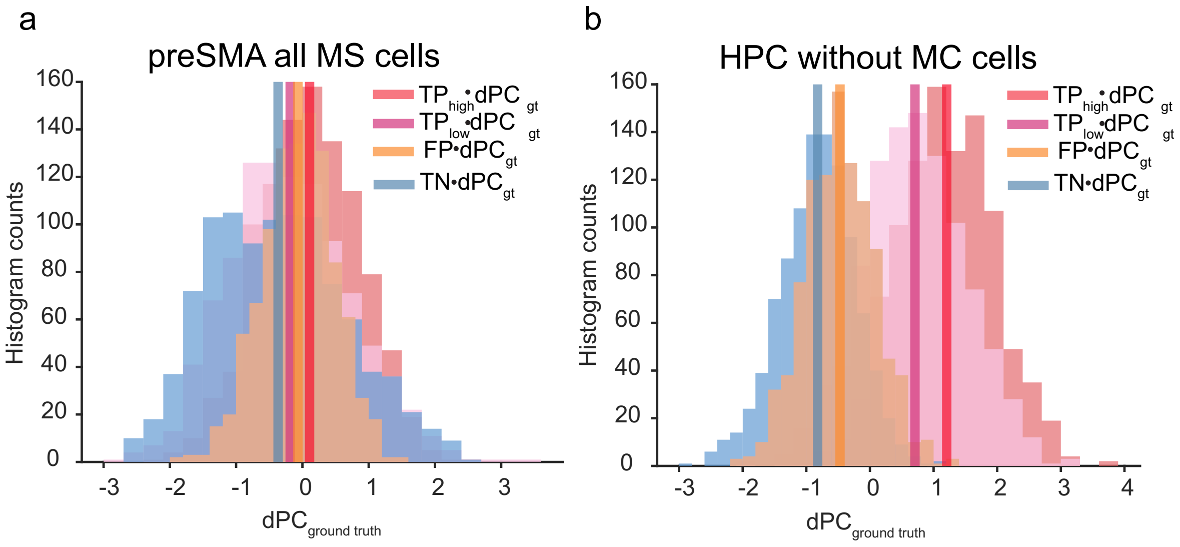


**Supplementary Figure 7. Additional dPCA projections analysis.**
(a-b) Distribution of single-trial projections onto dPC ground truth decoder axis for low-confidence TP, high-confidence TP, TN, and FP trials (pink, red, blue, and beige colors). In preSMA **(a)**, all trial types collapsed when projected onto the memory ground truth axis. In HPC, when MC cells were removed **(b)**, the high-confidence TP (red color) distribution was still the most separated from TN trials, with less separability for low-confidence TP and even less for FP trials, as when all MS cells are analyzed (Fig. 6f).

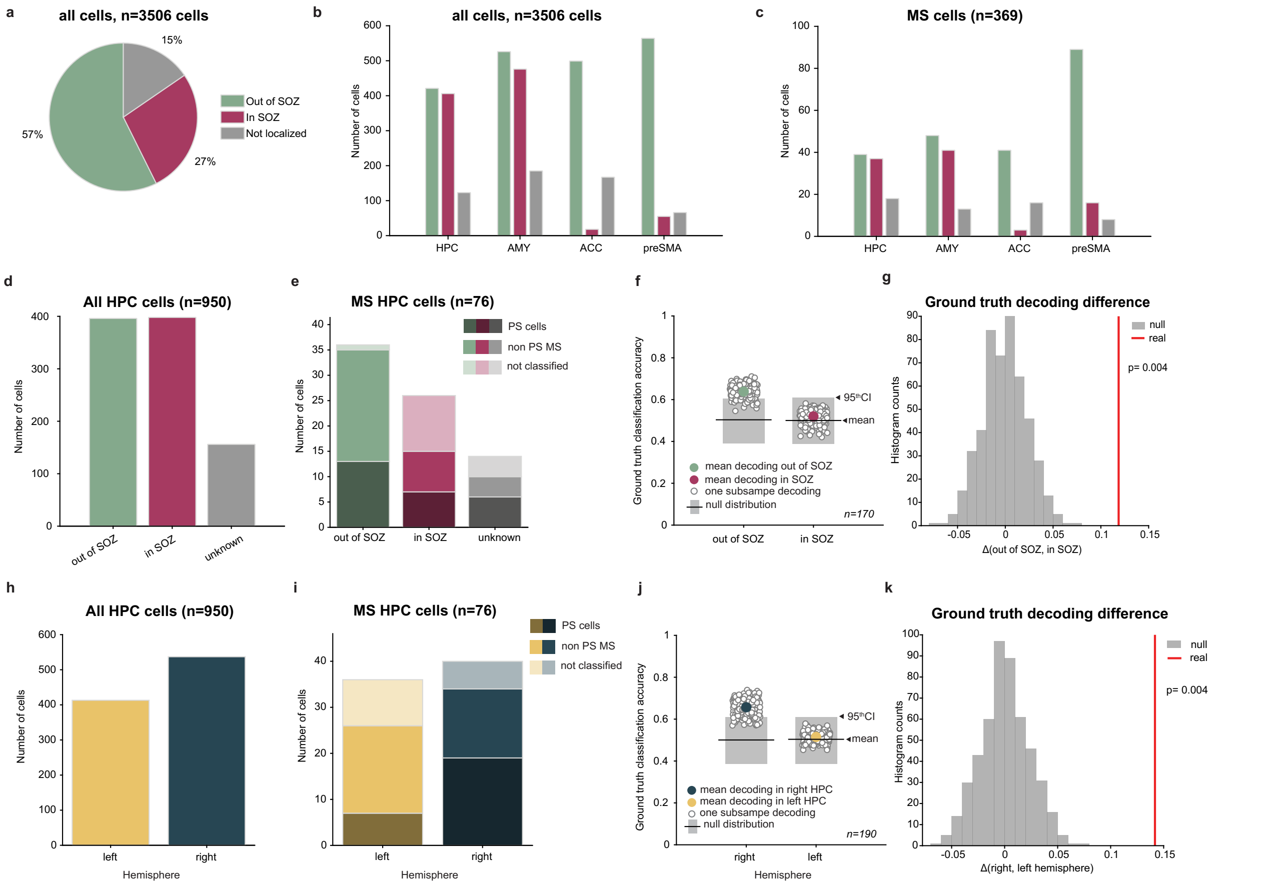


**Supplementary Figure 8. Epilepsy and laterality comparisons.**

**(a-c)** Distribution of cells inside or outside seizure onset zone (SOZ). **a-c,** Proportion of cells recorded that were located inside vs. outside the SOZ for all cells, for each brain area, and for only MS cells. **d,** Number of HPC cells recorded from within vs. outside the SOZ. **e,** Same as (d) but for analyzed hippocampal MS cells (excluding MS cells that are also tuned for visual categories). Different shades indicate if an MS cell belongs to PS type (dark shades) or not. **f,** Memory ground truth can be significantly decoded only from HPC cells that were recorded outside of the SOZ (accuracy=0.64, p= 0.019, in SOZ accuracy=0.52, p=0.39). For fair comparison, same number of cells (n=170, 85% of the smallest population) was randomly drawn from both in and out of SOZ population each decoder permutation (n=250). Colored dots indicate average across runs. **g,** Ground-truth decoding in out of SOZ HPC cells is significantly higher than in inside SOZ HPC cells (mean difference across n=250 subsamples, delta=0.12, red line). The null distribution (grey) is generated by shuffling the trial labels and repeating the same decoding procedure for both conditions and taking the difference. None of the permuted values exceeded the observed difference (p=0.004, p-value set as 1 divided by the number of permutations). **h,** Number of HPC cells recorded from the left (n= 413) and right (n=537) hemisphere**. i,** Same as (d) but for analyzed hippocampal MS cells (excluding MS cells that are also tuned for visual categories; n=36 and 40 for left and right respectively). Different shades indicate if MS cell belong to PS type (dark shades) or not. **j,** Memory ground truth can be significantly decoded only from HPC cells that were recorded from the right hemisphere (accuracy=0.66, p= 0.01, left accuracy=0.52, p=0.43). For fair comparison, same number of cells (n=190, 85% of the smallest population) was randomly drawn from both left and right population each decoder permutation (n=250). Colored dots indicate average across runs. **k,** Ground-truth decoding from the right hemisphere HPC cells is significantly higher than from the left (mean difference across n=250 subsamples, delta=0.14, red line). The null distribution (grey) is generated by shuffling the trial labels and repeating the same decoding procedure for both conditions and taking the difference. None of the permuted values exceeded the observed difference (p=0.004, p-value set as 1 divided by the number of permutations).
